# Self-balanced regulation by the long non-coding RNA *Lockd* on the cell cycle progression of cortical neural progenitor cells through counteracting *cis* and *trans* roles

**DOI:** 10.1101/2024.01.08.574564

**Authors:** Shaojun Qi, Jiangli Zheng, Qin Shen

## Abstract

Neural stem/progenitor cells (NSPCs) undergo active proliferation and exit the cell cycle upon precise regulation to produce differentiated progenies in order. Long non-coding RNAs (lncRNAs) have emerged as critical players in the developmental processes of NSPCs; however, relatively few have been shown to regulate the cell cycle *in vivo* directly. Here, we identified an NSPC-expressed lncRNA *Lockd* (lncRNA downstream of *Cdkn1b*) in the developing forebrain. Using *in vivo* loss of function models by premature termination of *Lockd* transcription via knockin polyadenylation signals or shRNA-mediated knockdown of *Lockd* (*Lockd*-KD), we show that *Lockd* is required for proper cell cycle progression of cortical NSPCs and the production of TBR2^+^ intermediate neural progenitor cells during cortical development. Interestingly, a comparison of genetic profiling in the two models reveals that *Lockd* promotes the expression of two counteracting cell cycle-related genes, *Cdkn1b in cis* and *Ccnd1 in trans*. Overexpression of *Ccnd1* or *Cdkn1b*-KD can rescue the cellular phenotypes of reduced cycling progenitors in *Lockd*-KD. Our results imply that lncRNA could act through distinct *cis* and *trans* mechanisms to achieve a self-balanced function.

## Introduction

During the development of the cerebral cortex, neural stem/progenitor cells (NSPCs) proliferate to establish and maintain a progenitor cell pool and also differentiate to generate neuronal progeny. Temporal and spatial mechanisms tightly control the cell-fate decision between proliferation and differentiation in NSPCs (Homem et al., 2015; Mitsuhashi and Takahashi, 2016; Telley et al., 2019). Specifically, in the early developing neocortex, NSPCs first symmetrically divide, expanding the progenitor cell population. These cells, first appearing as neuroepithelial cells, then radial glial cells (RGCs), are referred to as apical progenitor (AP) and located in the apical side of the ventricular zone (VZ). As development proceeds, APs can asymmetrically divide, contributing to neurogenesis directly or indirectly through generating the rapidly proliferating basal progenitors (BPs), also called intermediate progenitor cells (IPCs), which are the predominant neurogenic cell type at mid-corticogenesis (around embryonic day 14, E14 in mouse)(Farkas and Huttner, 2008; Haubensak et al., 2004; Noctor et al., 2004). It is known that cell cycle genes play an essential role in controlling neural cell proliferation. (Durak et al., 2016; Lange et al., 2009; Otsuki and Brand, 2018; Yoon et al., 2017). However, the mechanisms by which cell cycle genes are regulated in proliferating NSPCs are mostly unknown. (Cheffer et al., 2013).

The mammalian genomes are pervasively transcribed, producing tens of thousands of distinct non-coding RNAs that do not encode proteins (Consortium, 2012). Among them, long non-coding RNAs (lncRNAs), longer than 200 nucleotides (nt), have been shown to play essential roles in various biological processes, including neuronal differentiation (Andersen et al., 2019; Li et al., 2019b), X chromosome inactivation (Carmona et al., 2018; Engreitz et al., 2013; Jegu et al., 2017), inflammatory responses (Elling et al., 2018). Compared with protein-coding genes, lncRNAs are less evolutionarily conserved (Derrien et al., 2012). Remarkably, abundant lncRNAs are expressed in the brain, and most of them have temporally and spatially differential expression patterns, with some presenting highly restricted expression in only selected neuroanatomical structures or cell types (Aprea et al., 2013; Goff et al., 2015; Mercer et al., 2008). An increasing number of studies have investigated in mice the role of lncRNAs in regulating diverse essential biological processes during brain development, including mRNA stability (Faghihi et al., 2008), chromatin state (Lipovich et al., 2012; Liu et al., 2005; Modarresi et al., 2012), adjacent genes transcription (Bond et al., 2009; Feng et al., 2006) (Berghoff et al., 2013; Li et al., 2019b), mRNA splicing (Andersen et al., 2019; Ramos et al., 2015b). However, only a handful of lncRNAs have been functionally studied in the developing cerebral cortex *in vivo* (Andersen and Lim, 2018; Briggs et al., 2015; Wang et al., 2017). Moreover, the cellular and molecular mechanisms of lncRNAs underlying regulating cortical development are less clear.

The lncRNA locus can function *in cis* to influence the transcription of neighboring genes and/or *in trans*, to produce a molecule that leaves the site of transcription and affecting genes on a different chromosome or functions at distal cellular locations (Berghoff et al., 2013; Kopp and Mendell, 2018). Recent studies unveil that lncRNA can regulate the transcription of the adjacent transcription factor (TF) genes essential for NSPC fate choices (*in cis*)(Li et al., 2019b; Onoguchi et al., 2012). Additionally, in terms of *trans*functions, it has been reported that lncRNAs regulate the expression of genes required for cell fate specifications via directing TFs or chromatin modifiers to localize at their promoters (Chalei et al., 2014; Ng et al., 2013), or interacting with an RNA-splicing factor to regulate alternative splicing of transcripts essential for neural development (Andersen et al., 2019; Ramos et al., 2015a). Some lncRNAs function in both *cis* and *trans* manners to regulate multiple genes essential for the proliferation and neural differentiation of Neuro2a (N2A) cell lines (Chalei et al., 2014; Vance et al., 2014). However, for most lncRNAs, the *cis*-and/or *trans*-acting mechanisms and their relationship *in vivo* remain enigmatic (Bassett et al., 2014; Kopp and Mendell, 2018). Hence, in terms of the lncRNAs with both *cis* and *trans* effects, dissecting the contribution of *cis* versus *trans* mechanism to the phenotype *in vivo* can greatly improve our understanding of the biological significance of lncRNAs.

*Lockd*, a 434-nucleotide (nt) polyadenylated lncRNA, is abundant in many tissues, including the central nervous system. The deletion of the 25-kb *Lockd* gene locus reduces approximately 70% of the mRNA level of the upstream *Cdkn1b* (also known as p27kip1) gene, which is involved in the progression of cell cycle exit. (Paralkar et al., 2016; Tarui et al., 2005). However, premature termination of the *Lockd* transcription through the insertion of poly-adenylation (polyA) signals at 80 bp downstream of the *Lockd* transcription start site does not affect *Cdkn1b* transcription, suggesting the possibility of an enhancer-like *cis* element lying in the promoter region of *Lockd*, which is further confirmed by chromosomal conformation capture studies (Paralkar et al., 2016). In the nervous system, *Lockd* is abundant in areas such as the cerebral cortex (Paralkar et al., 2016), but its biological activity during cortical development is unknown.

Here, we report the expression pattern and function of *Lockd* in the developing cerebral cortex. We have identified the roles of *Lockd* using the shRNA knockdown and PolyA (poly-adenylation) knock-in approaches. Our results lead to a potential mechanistic model in which the *cis* and *trans* functions of *Lockd* contribute to balancing the cell cycle progression and neuronal differentiation of cortical NSPCs.

## Results

### The lncRNA *Lockd* is expressed in NSPCs but not in neurons during embryonic cortical development

We previously performed RNA sequencing (RNA-seq) analysis of cortical NSPCs from the Ebf2-EGFP mouse embryos (Figure 1A)(Chuang et al., 2011; Li et al., 2019a). Among the genes that are differentially expressed in E11.5 cortical NSPCs (with the expression level of FPKM > 1 and fold change > 2 compared with that in the preplate neurons), we identified 25 NSPC-enriched lncRNAs (Figure 1A). We focused on one of the top 5 most abundant lncRNAs in the E11.5 NSPCs, *Lockd* (lncRNA downstream of *Cdkn1b*), as its function in the nervous system has not been studied before.

**Figure 1.**
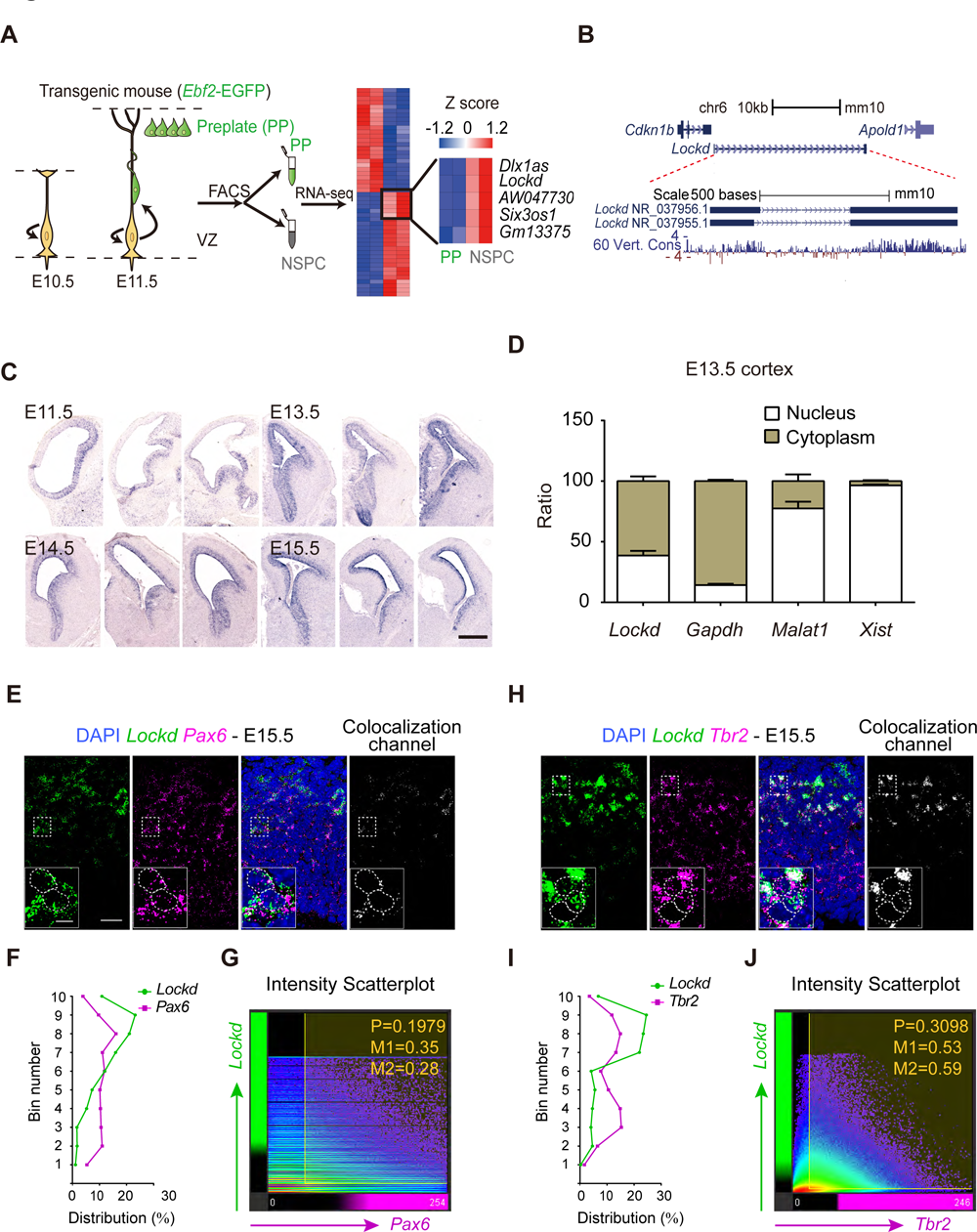
Characterization of *Lockd*, a lncRNA Enriched in NSPCs. (A) Schematic explaining the steps of lncRNAs selection. E11.5 Ebf2-EGFP transgenic mouse brains were used to isolate the GFP negative NSPCs by Fluorescence-activated Cell Sorting (FACS). RNA sequencing (RNA-seq) analysis shows the NSPCs enriched lncRNAs, and several items are indicated in an enlarged heatmap. (B) UCSC genome browser view of *Lockd* locus. Tracks depicting mammalian conservation (Placental Cons), as well as NCBI Ref-seq genes, are shown. (C) *In situ* hybridization (ISH) of *Lockd* in embryonic mouse brains from different stages. Scale bar, 500 μm. (D) Subcellular fractionation of indicated lncRNAs and mRNAs in E13.5 cortex tissue culture followed by RT-qPCR. Error bars depict mean ± SEM from three biological replicates. (E-J) Shown in (E) and (I) are representative images of SABER-FISH detection for *Pax6* or Tbr2 and *Lockd* in E15.5 brain sections. Scale bar, 20 μm. Insets display the subcellular localization of *Lockd* transcripts at higher magnification, and the dashed outlines indicate the boundary of a nucleus. Scale bar, 5 μm. Shown in (F) and (I) are quantifications for the percentage of SABER-FISH dots in each bin within 160 μm distance from the apical ventricular surface. A Scatter plot of signal intensities for *Lockd* versus *Pax6* (G) or *Tbr2* (J) was generated by Imaris software. Colocalization was represented as a new channel corresponding to the colocalized voxels (E and H). The Pearson coefficient for colocalized voxels and Manders colocalization coefficients (M1 and M2) are indicated in the graphs (G and J).

The *Lockd* is encoded on chromosome 6 and has two isoforms annotated in the NCBI Refseq database (Figure 1B), with NR_037956.1 as the main isoform in NSPC (FPKM of NR_037956.1 = 4.44, FPKM of NR_037955.1 = 1.58). Analysis of RNA-seq datasets from NCBI indicates that *Lockd* is highly expressed in the developing brain (Figure S1A). We found by *in situ* hybridization (ISH) that *Lockd* is expressed explicitly in the ventricular and subventricular zone (VZ-SVZ) but not in the cortical plate of the mouse forebrain at different embryonic stages (Figure 1C). As the subcellular localization of a lncRNA in cells (such as nucleus, cytoplasm, or specific organelles) is closely related to its function (Leone and Santoro, 2016), we performed subcellular fractionation with cultured E13.5 cortical cells or the N2a cells followed by RT-qPCR analysis. *Lockd* was enriched in the cytoplasmic component while also present in the nuclear proportion (Figure 1D and S1B). To further determine the identities of *Lockd* expressing cells *in vivo*, we employed fluorescence *in situ* hybridization (FISH) assay with the SABER (signal amplification by exchange reaction) amplification technology applied to boost signal detection (Figure S1C). SABER has been proved to be efficient and straightforward to achieve highly amplified RNA FISH signals in fixed cells and tissues (Kishi et al., 2019). We used a pool of two primary FISH probes to simultaneously detect the transcripts of *Pax6*, a marker for radial glial cells, and *Lockd (Pax6*/*Lockd)*, or *Tbr2*, a marker of BPs, and *Lockd* (*Tbr2*/*Lockd*) in the E15.5 mouse forebrain (Figure 1E and 1H). The signals of *Lockd* were prominent in the VZ-SVZ region and enriched in *Pax6^+^* and *Tbr2^+^* cells, with an apical low-basal high gradient (Figure 1F and 1I). Consistent with the subcellular fractionation studies, we also observed that *Lockd* transcripts predominantly localized in the cytoplasm of NSPCs (Figure 1E and 1H). Notably, *Lockd* colocalized more with *Tbr2* than *Pax6 in vivo* (Figure 1G and 1J).

### Early termination of the nascent *Lockd* transcript leads to accelerated cell-cycle progression and premature neuronal differentiation of NSPCs

Prior *in vitro* studies in mouse erythroid cell line G1E have identified the promoter of *Lockd* contains an enhancer-like *cis-*element, which regulates the upstream gene *Cdkn1b* by associating with its promoter (Paralkar et al., 2016). To determine whether the promoter or exon1 region of *Lockd*harbors any enhancer marks in the nervous system, we analyzed the chromatin state using histone modification profiles from the Encyclopedia of DNA Elements (ENCODE) project datasets of E13.5 mouse forebrains (Gorkin et al., 2017). The analysis included histone 3 lysine 4 mono-methylation (H3K4me1), a chromatin mark enriched in poised and active enhancers, and histone 3 lysine 4 tri-methylation (H3K4me3), an epigenetic modification associated with promoter regions, as well as acetylation of lysine 27 of histone 3 (H3K27ac), a well-accepted active mark of promoters or enhancers (Heintzman et al., 2009; Heintzman et al., 2007). All tracks were integrated and displayed with the UCSC genome browser. We found a relatively high level of H3K4me3-over H3K4me1-modification in the regions of *Lockd*promoter and exon1, which were also enriched in H3K27ac, and these regions were annotated with an active promoter-like chromatin state. In contrast, enhancer marks (H3K4me1 ^high^ H3K4me3^low^) were identified at an upstream intergenic region of *Lockd* (Figure 2A). Hence, *Lockd* likely belongs to promoter-associated lncRNAs (plncRNAs), a class of lncRNAs produced from promoter-like (H3K4me3^high^) transcriptional initiation regions (Marques et al., 2013). These data suggest no enhancer activity lying within exon1 of *Lockd*, which is consistent with the previous findings (Paralkar et al., 2016).

**Figure 2.**
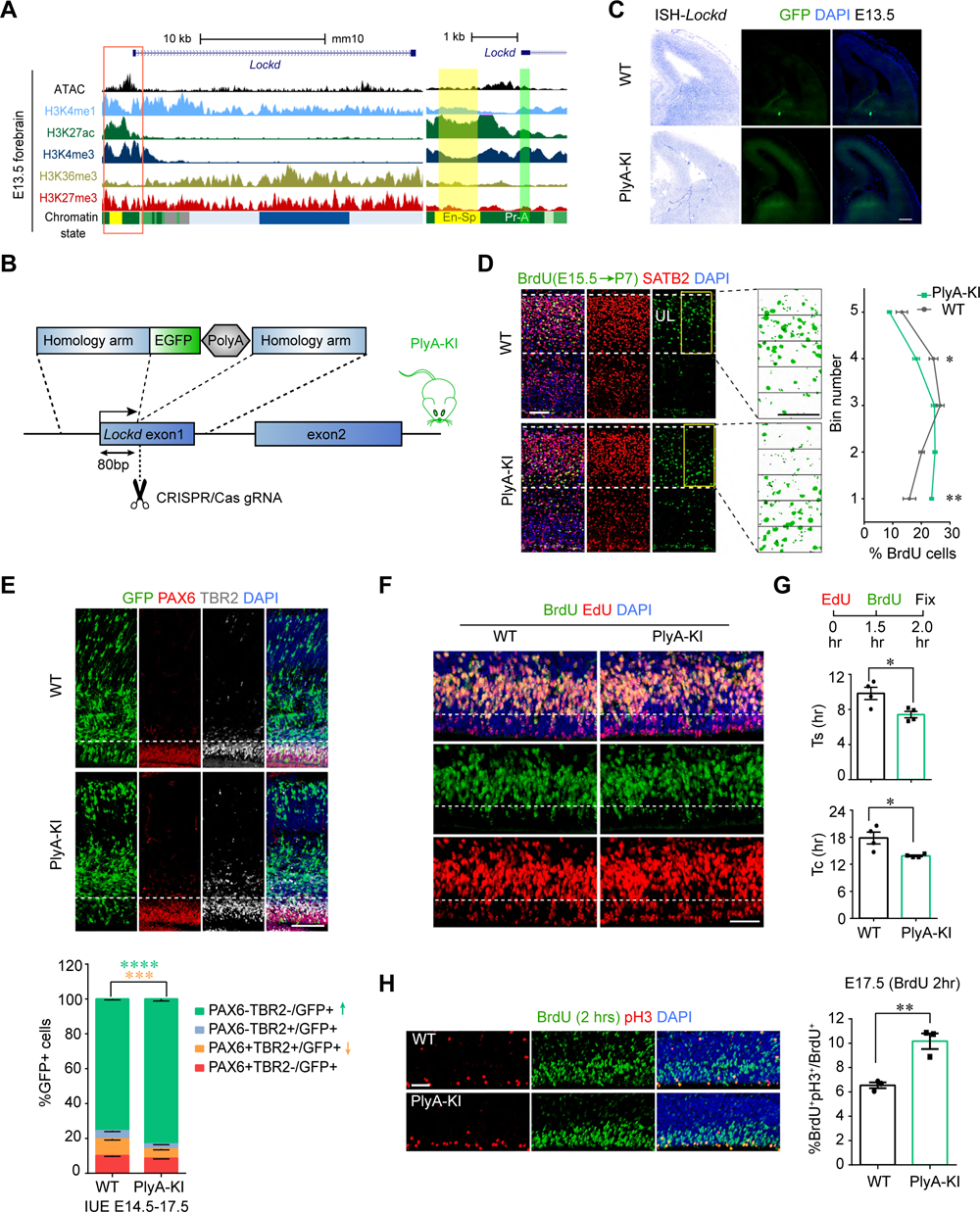
Poly-A insertion in *Lockd* locus results in a reduction of the transitioning NSPCs and premature neuronal differentiation. (A) Histone modifications and chromatin state profile of *Lockd* gene locus. Inset shows a closer view of upstream and near the transcription start site of *Lockd* (right). Promoter, Active (Pr-A); Enhancer, Strong TSS-proximal (En-Sp). (B) Schematic of *Lockd* truncated mouse line used. CRISPR/Cas9-induced homologous recombination locates 80bp downstream of the *Lockd* transcription start site, inserting an EGFP-PolyA (poly-adenylation) cassette to terminate the nascent *Lockd*transcript. (C) ISH of *Lockd* (left) and GFP immunostaining (right) on coronal sections of E13.5 mouse cortex from WT (top) and PlyA-KI (bottom). Scale bar, 200 μm. (D) E15.5 labeling with BrdU followed by the analysis of cell type and distribution at P7. Left: sample confocal images stained with SATB2, BrdU, DAPI. The area between white dashed lines is the UL with dense SATB2 staining. The area indicated by the yellow box is divided into five bins on average. Scale bars, 100μm. The proportion of BrdU^+^ neurons in each bin was quantified (right) (n = 3). The data represent the mean ± SEM. The statistical analysis is two-way ANOVA, followed by Sidak’s multiple comparison test (*P < 0.05; **P < 0.01). (E) Immunohistochemistry (IHC) for PAX6, TBR2 9. After E14.5 *in utero* electroporation (IUE), the fractions of PAX6^+^/TBR2^−^, PAX6^+^/TBR2^+^, TBR2^+^/PAX6^−^ and PAX6^−^/TBR2^−^ in total GFP^+^ cells were quantified at E17.5 (n = 4,WT; n = 3, PlyA-KI). Data represent mean ± SEM; statistical analysis is two-way ANOVA followed by Sidak’s multiple comparisons tests (***p < 0.001, ****p < 0.0001). Scale bar, 100 μm. (F and G) Analysis for cell-cycle duration. Pregnant mice were injected with EdU, followed by BrdU 1.5 hrs later, and sacrificed at 2 hrs. The WT and PlyA-KI cortical sections were immunostained for BrdU and EdU (F, scale bar, 50μm). Cell cycle kinetics for (G, up) mean S-phase duration (Ts), (G, lower) mean total cell cycle duration (Tc) were quantified. Values represent mean ± SEM (n = 4, WT; n = 4, PlyA-KI; *p < 0.05; unpaired Student’s t test). (H) S to M phase transition analysis. Two hours after injection with BrdU at E17.5, the WT and PlyA-KI cortical sections were immunostained for BrdU and pH3. (left, scale bar, 50μm). Right: the fraction of pH3/BrdU^+^ cells relative to BrdU^+^ cells was quantified (n = 5, WT; n = 6, PlyA-KI). Values represent mean ± SEM (*p < 0.05; unpaired Student’s t test).

To investigate the biological function of *Lockd in vivo* without eliminating the *cis-*element of the promoter region, we generated a mouse model to block the full-length transcription of *Lockd* (animals carrying an EGFP-PolyA cassette inserted by clustered regularly interspaced short palindromic repeats (CRISPR)-Cas9-mediated homologous recombination at 80bp downstream of *Lockd* gene transcription start site)(Figure 2B). In the polyA cassette knock-in mouse (referred to as PlyA-KI), *Lockd* expression in the VZ-SVZ was ablated as assessed by ISH (Figure 2C). Additionally, EGFP expression driven by the endogenous *Lockd* promoter revealed cells residing in the VZ-SVZ, consistent with the ISH pattern (Figure 2C). However, the EGFP signal was too weak to be detected (3 slides from four mice examined) when we switched from wide-field fluorescent microscopy to confocal microscopy (Figure S2A). Nevertheless, the absence of the EGFP signal from the PlyA-KI mice allowed us to use the GFP channel for detecting electroporated cells and staining in the following experiments.

The development of mouse cortical layers occurs in a time-dependent manner: NSPCs first produce projection neurons residing in the deeper layers (L5 and L6), and later-born neurons migrate through these early-generated neurons to form the upper layer (L2/L3 and L4)(Lodato and Arlotta, 2015). The production of the upper layer neurons peaks around E15.5, when PAX6-positive RGCs produce TBR2-positive IPs, which further differentiate into SATB2-positive neurons (Taverna et al., 2014; Vasistha et al., 2015). To study the effects of PlyA-KI on cortical development, we intraperitoneally injected pregnant mice with BrdU at E15.5. We then examined the neuronal subtypes and cortical distributions of BrdU^+^ cells at postnatal day (P) 7. Consistently, we found that nearly all BrdU^+^ cells in the cortical plate were SATB2-positive neurons in both WT and PlyA-KI mice (data not shown), indicating that the cell fate of BrdU labeled cells at E15.5 in PlyA-KI mice was not changed. However, although BrdU^+^ cells in PlyA-KI mice migrated into the upper layer (UL), the location of BrdU^+^ cells was relatively deeper compared with that of WT (Figure 2D). These results suggest that E15.5 *Lockd*-deficient NSPCs differentiate into neurons earlier than WT since the birthdate determines their final location in the cortex.

Given the different distribution of BrdU^+^ neurons at P7, we assessed the proliferation and differentiation of NSPCs at the embryonic stages. To characterize the impact of the loss of *Lockd* transcripts on cortical NSPC lineage progression, we electroporated a control plasmid expressing GFP into the cortex of the WT and PlyA-KI embryos at E14.5 and analyzed GFP^+^ cells at E17.5. Compared to that of WT controls, there was a significant decrease in the percentage of PAX6/TBR2 co-expressing GFP^+^ cells (RGs transitioning into IPs) accompanied by more GFP^+^ cells being PAX6^-^/TBR2^-^ (immature neurons) in the PlyA-KI cortices (Figure 2E), indicating that PlyA-KI leads to the premature differentiation of NSPCs in the developing cortex. We also quantified the total number of PAX6^+^ and TBR2^+^ cells in the E15.5 cortex, and the percentage of PAX6^+^/TBR2^+^ transitioning cells among total TBR2^+^ IPCs was consistently reduced in PlyA-KI mice (Figure S2B).

Previous studies have shown that cell cycle progression plays an essential role in the process of asymmetric division of RGC to form IP and then to differentiate into neurons (Furutachi et al., 2015; Pilaz et al., 2009). The reduction of PAX6^+^/TBR2^+^ cells in PlyA-KI mice could be due to the defects in cell cycle progression. To test this, we performed the dual-labeling thymidine analog approach by consecutive injections of EdU and BrdU at E14.5 to determine the mean S-phase duration (Ts) and total cell cycle duration (Tc) within NSPCs (Figure 2F and 2G). We found that the NSPCs in PlyA-KI mice had a significantly shorter S-phase (WT= 9.83 ± 0.70 hrs; PlyA-KI= 7.41 ± 0.37 hrs) and total cell cycle duration (WT= 17.83 ± 1.31 hrs; PlyA-KI= 13.87 ± 0.18 hrs) than WT NSPCs (Figure 2G). We also stained for the M phase marker (phospho-histone 3, pH3) at 2 hrs after BrdU pulse labeling to directly examine the S to M phase at a later stage (E17.5)(Figure 2H). There was a significant increase in the ratio of BrdU^+^/pH3^+^ cells among all BrdU^+^ cells in PlyA-KI mice, indicating more cells transited from S to M phase during the 2-hr chase (Figure 2H). To further examine the cell-cycle exit of proliferating NSPCs, we labeled cycling NSPCs with BrdU at E16.5 and analyzed the expression of Ki67 (a proliferation marker) at E17.5. The fraction of cells exiting the cell cycle (BrdU^+^Ki67^−^cells / BrdU^+^ cells) was increased in the PlyA-KI mice (Figure S2C and S2D). To quantify cell-cycle progression at the population level, we pulsed cortical neural progenitor cells (NPCs) derived from E13.5 mouse cortex with EdU for 30 mins and cultured cells for 5 hrs, followed by the flow cytometry analysis for the cell-cycle status of NSPCs (Figure S2E). We found a significant increase in the percentage of EdU^+^ NSPCs divided after 5 hrs within the PlyA-KI group compared to the WT control, indicating an accelerated cell cycle progression (Figure S2F). Together, these results suggest that *Lockd* is required to regulate the cell cycle progression of NSPCs during forebrain development.

### The expression of *Lockd* is essential in the cell cycle progression and has effects on its neighboring genes *in cis*

To identify the changes in gene expression profile caused by loss of *Lockd* transcript, we performed RNA-seq of the mouse cortex tissue at E13.5, a stage enriched for NSPCs (Figure 3A). Differential gene expression analysis revealed 141 downregulated genes and 276 up-regulated genes between WT and PlyA-KI cortex (P < 0.05, fold change > 1.2, criteria used previously (Durak et al., 2016)). A subset of differentially expressed genes (DEG) was further confirmed via qPCR (Figure 3B). After filtering with the cell-type-specific gene pools collected from the datasets (Telley et al., 2019), we found that most downregulated genes (29 of 34, 85.3%) belonged to the radial glia (RG)-like gene set, sharing a similar expression pattern with *Pax6* (also include *Pax6*^+^, *Tbr2*^+^ transitioning cells); The majority of up-regulated genes (55 of 62, 88.7%) were grouped in the Neuron-like gene population (N), showing expression pattern similar to *Neurod4*. In contrast, much fewer DEGs displayed intermediate progenitor (IP)-like enrichment such as *Tbr2* (Figure 3C), indicating that *Lockd* regulates stage-specific transcriptional programs.

**Figure 3.**
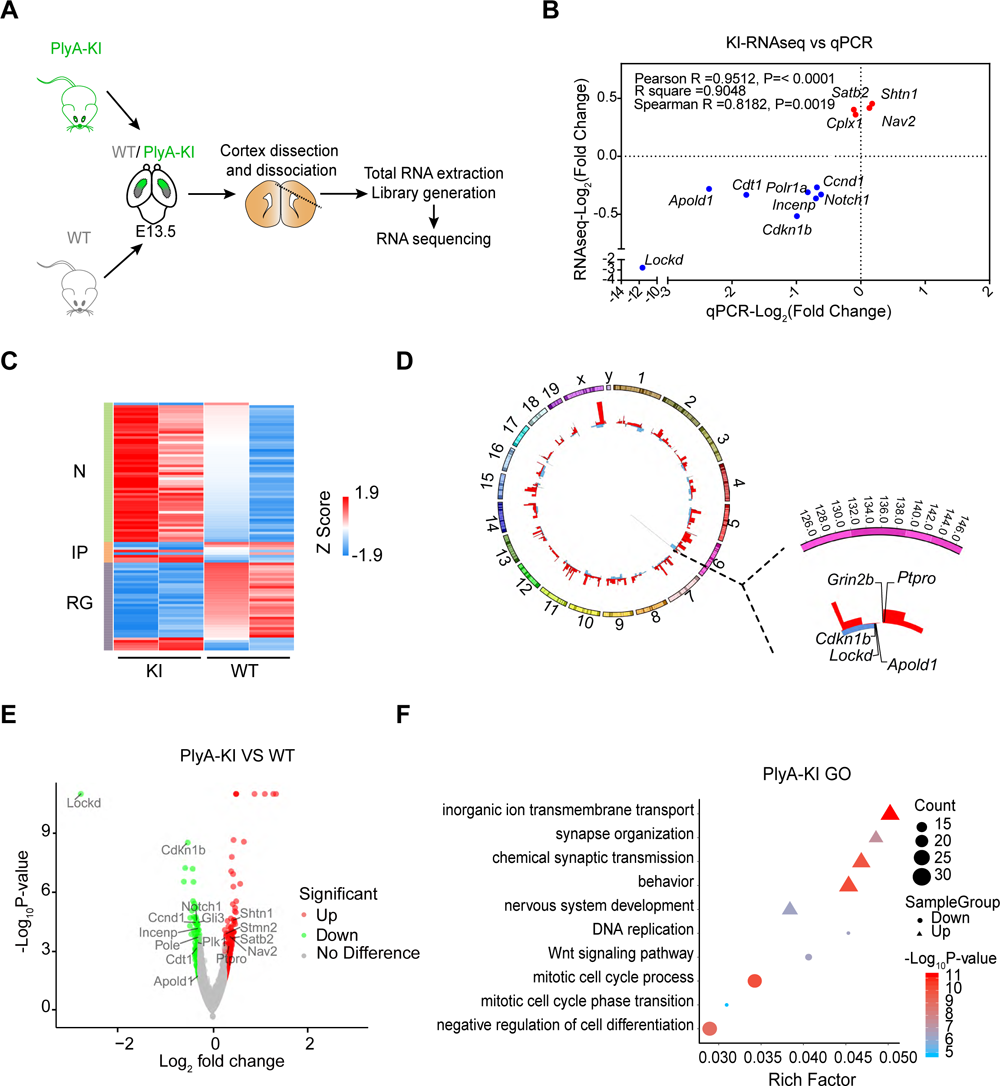
*Lockd* is involved in the regulation of cell cycle progression and have *cis* effects on its neighboring genes. (A) Schematic for transcriptional profiling of the WT or PlyA-KI mouse cortex at E13.5. (B) The mRNA level confirmation of several candidate DEGs identified by RNA-Seq using qRT-PCR for the E13.5 cortex cells normalized to *Gapdh.* The correlation coefficient was computed between the log_2_ fold change of RNA-seq and qRT-PCR values. (C) Heatmap represents the differentially expressed genes (DEGs) in WT, PlyA-KI biological duplicate samples. Cluster analysis displays the enrichment of DEGs in radial glia (RG), intermediate progenitors (IP), and neurons (N). (D) Circos plot of genomic locations of DEGs upon *Lockd* PlyA-KI. The expanded region displays the adjacent DEGs of *Lockd* within 2 Mb. (E) The volcano plot shows the fold change (> 1.2 fold) in expression and its significance (p < 0.05), with each dot representing a gene. (F) Gene ontology analysis of DEGs between WT and PlyA-KI RNA-seq data.

LncRNAs can either control the expression of their neighboring genes *in cis* or leave the site of their transcription and affect genes on different chromosomes or mRNAs located at cytoplasm in a *trans* manner (Elling et al., 2018; Kopp and Mendell, 2018; Yin et al., 2015). Thus, we analyzed the location of the DEGs and found that PlyA-KI altered the expression of genes the closest to *Lockd*in the genome (± 2.0Mb from *Lockd*), including *Cdkn1b*, *Apold1,* and also genes within 2.5Mb (*Grin2b* and *Ptpro*)(Figure 3D), suggesting that *Lockd* regulates its neighboring genes *in cis* via a lncRNA transcript-mediated mechanism. Interestingly, among these *Lockd* neighboring genes, *Cdkn1b* is the most significantly down-regulated (Figure 3E).

To identify the biological functions affected by the truncated *Lockd*, we performed gene ontology (GO) analysis of the differentially expressed genes. We found that the downregulated genes were mostly involved in regulating the cell cycle, and the up-regulated genes were enriched for the processes relating to mature neuronal function (i.e., synapse organization and synaptic transmission) (Figure 3F). Among the down-regulated genes, those in the cell cycle-related GO terms, such as the mitotic cell cycle process and cell cycle phase transition, were significantly enriched (Figure 3F). Surprisingly, in addition to *Cdkn1b*, *Cyclins D1* (*Ccnd1*), the critical component of cell cycle machinery with an opposite role to *Cdkn1b*, was significantly repressed in the PlyA-KI mouse (Figure 3B and 3F), suggesting *Lockd* regulates cell cycle genes with different mechanisms.

### The *Lockd* RNA regulates neural proliferation and differentiation via *trans* **manner**

The short hairpin RNA (shRNA) strategy for knocking-down (KD) the expression of *lncRNAs* allows the least disturbance to their *cis* functions on the neighboring genes. It is particularly effective for testing the *trans*-function of cytoplasmic lncRNAs as shRNAs are processed to silence the cytoplasm targets and have a much lower or no efficiency in the nuclear fraction (Liang et al., 2011; Lima et al., 2016). We electroporated the *Lockd*-KD (sh*Lockd*-1-RFP) or control (shCtrl-RFP) construct into the VZ cells at E14.5 (Figure 4A). N2a cells transfected with the same shRNA (sh*Lockd*-1) led to an efficient knockdown of endogenous *Lockd* transcripts (Figure S3B). Two days after *Lockd*-KD, the proportion of transitioning cell types from RGs to IPs (PAX6^+^/TBR2^+^/RFP^+^ cells) was reduced, while the proportion of PAX6^-^/TBR2^-^/RFP^+^ young neurons increased (Figure 4A). We also co-electroporated the constructs of sh*Lockd*-1-RFP or shCtrl-RFP together with *pNeurod1*-EGFP, an indicator for neurogenic cells (GFP^+^/RFP^+^ cells) at E14.5 to evaluate the effect of knocking-down of *Lockd* on neurogenesis two days later (Fang et al., 2013; Guerrier et al., 2009; Heng et al., 2008). Compared to the control group, more RFP^+^ cells in the *Lockd*-KD were *pNeurod1*-EGFP^+^ and appeared out of the SVZ (Figure S3C).

**Figure 4.**
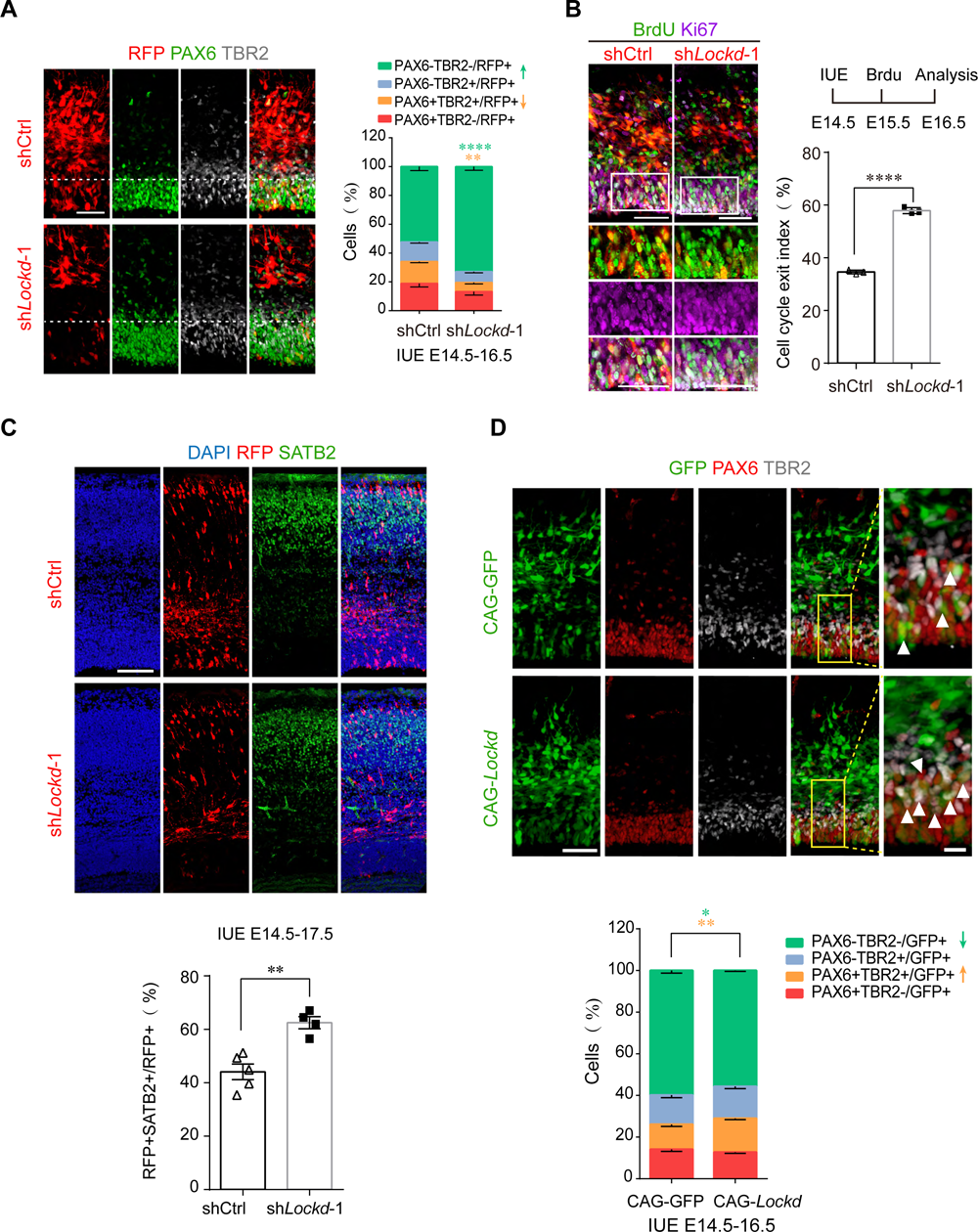
*Lockd* regulates the differentiation of mouse cortical progenitors *in trans*. (A) Analysis of neural differentiation after knockdown (KD) of *Lockd*. IUE at E14.5, the fractions of PAX6^+^/TBR2^-^, PAX6^+^/TBR2^+^, TBR2^+^/PAX6^-^ and PAX6^-^/TBR2^-^ in total RFP^+^ cells were quantified at E16.5 (n = 6, shCtrl; n = 5, sh*Lockd*-1). Data represent mean ± SEM; statistical analysis is two-way ANOVA followed by Sidak’s multiple comparisons tests (**p < 0.01, ****p < 0.0001). Scale bar, 50μm. (B) Images of Ki67 (purple) and BrdU (green) staining in the E16.5 cortices electroporated with shCtrl, sh*Lockd*-1 at E14.5. BrdU was labeled 24h before brain dissection. The white box indicates the region expanded in the lower panels. Scale bars, 50μm. Quantification of Ki67^-^/Brdu^+^/RFP^+^ cells relative to Brdu^+^/RFP^+^ cells was calculated as the cell cycle exit index. Values represent mean mean ± SEM (n=3 ; ****p < 0.0001; unpaired Student’s t test). (C) Cortical sections of E17.5 electroporated at E14.5. Images were stained for SATB2 (green), DAPI (blue). Scale bar, 100 μm. The lower plot shows the quantification of SATB2^+^/RFP^+^ as a percentage of total RFP^+^ cells. Values represent mean ± SEM (n=5 for shCtrl and n=4 for sh*Lockd*-1; **p < 0.01; unpaired Student’s t test). (D) Mouse cortices were electroporated at E14.5 to express GFP (control: CAG-GFP) and *Lockd* (CAG-*Lockd*) and analyzed at E16.5. Scale bar, 50 μm. The regions of white boxes are enlarged in adjacent panels. Arrowheads point to PAX6^+^/TBR2^+^/ GFP^+^ cells. Scale bar, 10 μm. The percentage of PAX6^+^/TBR2^-^, PAX6^+^/TBR2^+^, TBR2^+^/PAX6^-^ and PAX6^-^/TBR2^-^ among total GFP^+^ cells were quantified as a lower panel. Values represent mean ± SEM (n=4 for CAG-GFP and n=5 for CAG-*Lockd*; *p < 0.05; **p < 0.01; two-way ANOVA followed by Sidak’s multiple comparisons test).

We further labeled cycling progenitor cells with BrdU at E15.5 and analyzed the fraction of BrdU-labeled cells co-stained with Ki67 24 hr later. We found that *Lockd*-KD reduced the number of BrdU^+^/Ki67^+^ cells, whereas it increased the number of BrdU^+^/ Ki67^-^ cells, leading to an increased cell cycle exit index (the ratio of BrdU^+^Ki67^-^/BrdU^+^)(Figure 4B), indicating silencing *Lockd* leads to accelerated differentiation of NSPCs. Consistently, we found that *Lockd*-KD resulted in more SATB2^+^ neurons in a longer time window of three days after electroporation (Figure 4C).

To further assess the function of *Lockd* in producing PAX6^+^/TBR2^+^ transitioning IPs, we overexpressed *Lockd* in RGs at E14.5 by electroporating CAG-*Lockd* or control-construct CAG-GFP and followed by a cell type analysis of the electroporated cells. There was a significant increase in the fraction of PAX6^+^/TBR2^+^/GFP^+^ IPs and a mild increase of young neurons (PAX6^-^ /TBR2^-^/GFP^+^) in brains electroporated with CAG-*Lockd* compared to control (Figure 4D). Together, these results indicate that the *trans* action of *Lockd* regulates the proliferation and differentiation of neural progenitors and is required to transition from RGs to IPs then to neurons.

### *Lockd* functions *in trans* to regulate a core network of the cell cycle genes

To further understand the molecular mechanism of *Lockd trans* function during neural development, we explored the changes in the transcriptome of NSPC lineages following *Lockd*-KD. We electroporated shCtrl-GFP or sh*Lockd-*1-GFP at E14.5 and isolated these GFP^+^ cells with fluorescence-activated cell sorting (FACS) at E16.5, followed by RNA-seq for transcriptional profiling (Figure 5A). Differential gene expression analysis revealed 693 genes in sh*Lockd* samples, which were differentially expressed compared with shCtrl groups (P < 0.05, fold change > 1.2). Similar to the changes caused by loss of *Lockd* in the PolyA-KI mice, most down-regulated genes were enriched in the RG-like gene set, while only a handful of down-regulated genes displayed IPs enrichment. Moreover, the majority of up-regulated genes were grouped in the Neuron population (Figure 5B).

**Figure 5.**
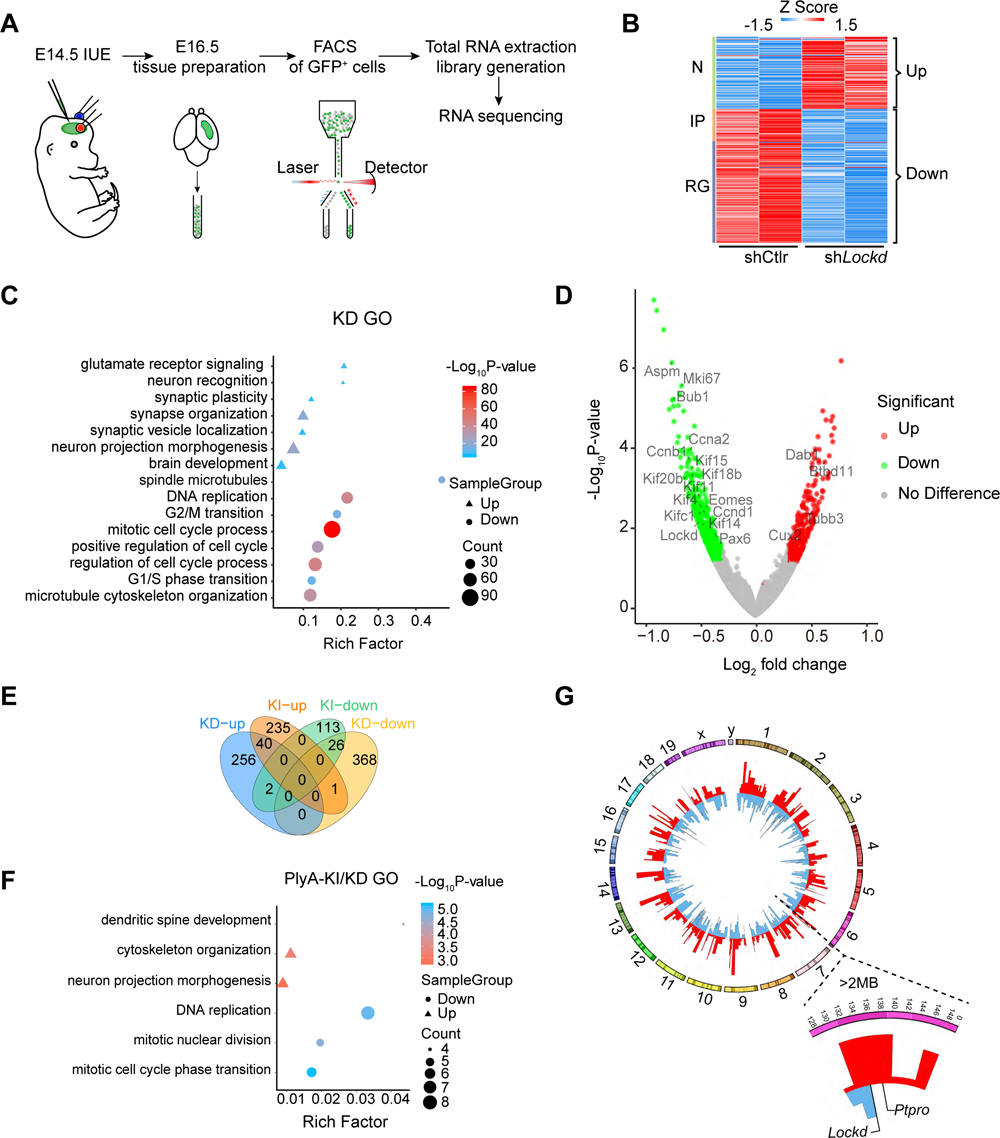
*Trans* roles of *Lockd* regulate a core network of genes linking to the cell cycle. (A) Schematic illustration of RNA-seq assay. (B) Heatmap shows the differentially expressed genes (DEGs) and cluster analysis for the enrichment of DEGs in radial glia (RG), intermediate progenitors (IP), and neurons (N). (C) Analysis of significant genes ontology of biological process (GO) for DEGs from *Lockd*-knockdown RNA-seq data. (D) The volcano plot illustrates the fold change (> 1.2 fold) in expression and its significance (p < 0.05), with each dot representing a gene. Several genes involved in the progression of the cell cycle or neural differentiation are depicted within the graph. (E) Venn diagrams display the DEGs overlapped between PlyA-KI and KD. The number of genes is indicated. (F) Analysis of significant genes GO terms for overlapped DEGs from PlyA-KI and KD RNA-seq data. (G) Circos plot shows the genomic locations of DEGs upon *Lockd* knockdown. The expanded region displays the adjacent DEGs of *Lockd* within 2 Mb.

Next, GO analysis of *Lockd*-KD showed a similar enrichment of biological processes in PlyA-KI: downregulated genes were mostly involved in the cell cycle progression, and up-regulated genes were enriched for neuronal maturation (i.e., synapse organization and neuron morphogenesis)(Figure 5C). For example, cell cycle-related genes such as *mKi67*, *Aspm*, *Ccnd1* were downregulated while neuronal genes such as *Satb2*, *Tubb3* were up-regulated (Figure 5D). We further verified these changes with RT-qPCR (Figure S4A). By comparing the DEGs of PlyA-KI and KD models, there were 26 common down-regulated genes and 40 up-regulated genes in both models (Figure 5E). GO analysis revealed the overlapped down-regulated genes were involved in cell cycle progression, and both up-regulated genes had reported roles in neuronal maturation (i.e., neuron projection morphogenesis)(Figure 5F). We also analyzed the location of DEGs on the genome; however, we found that *Lockd*-KD did not affect the expression of its nearby transcripts (± 2.0Mb from *Lockd*, including *Cdkn1b* and *Apold1*)(Figure 5G), confirming the regulation on *Cdkn1b* by *Lockd* is likely due to the *cis*action but not the *trans* action.

To further confirm the *trans* functions of *Lockd*, we overexpressed *Lockd* in RGs by IUE with CAG-*Lockd*-GFP at E14.5 and sorted out these GFP^+^ cells by FACS at E16.5, followed by RT-qPCR detection (Figure S4B). Overexpression of lncRNA *Lockd* did not affect the expression of *Cdkn1b* (closest gene to *Lockd*) but increased the mRNA level of *Ccnd1* (Figure S4B).

### The distinct *cis and trans* functions of *Lockd* are required to balance neural proliferation and differentiation

As the PlyA-KI mediated truncation of the *Lockd* transcript disrupts both *cis* and *trans* functions of the *Lockd* RNA transcript, while the shRNA mediated *Lockd* KD only affects the *trans* manner, we were able to further dissect the *cis* and *trans* functions of *Lockd* with the two different test models. Interestingly, we noticed both cellular and molecular phenotypic difference comparing the two models. Although both PlyA-KI and *Lockd*-KD led to defective NSPC proliferation and differentiation, the effect in *Lockd*-KD was severer than that in PlyA-KI. For example, the reduction in the proportion of PAX6^+^/TBR2^+^ cells was more than twofold in *Lockd*-KD compared to KI (WT-KI = 4.30%; shCtrl – KD = 8.97%) (Figure 2E and 4A). Also, we found that the *Lockd* neighboring gene *Cdkn1b* was only downregulated in the PlyA-KI model (Figure 3D and 3E), while *Ccnd1* was downregulated in both models (Figure 3B, 3E, and 5D). Consistently, we found that CCND1 protein was also significantly reduced in PlyA-KI mice by immunostaining (Figure S5A).

*Cdkn1b* is one of the major cyclin-dependent kinases (CDK) inhibitors (CKIs) required for suppression of CDK activity (Massague, 2004; Sherr and Roberts, 1999) and is involved in the progress of cell cycle exit in cortical neural precursors (Goto et al., 2004; Kiyokawa et al., 1996; Tarui et al., 2005). On the other hand, *Ccnd1* is expressed from the mid-to-late-G1 phase and controls G1 lengthening to regulate progenitor expansion or neurogenesis (Lange et al., 2009; Wong et al., 2015). *Cyclin D1*-deficient mice (*Ccnd1*^-/-^) display developmental abnormalities, hypoplastic retinas, and pregnancy-insensitive mammary glands (Fantl et al., 1995; Sicinski et al., 1995).

To investigate whether *Lockd* regulates the proliferation and differentiation of NSPCs through *Ccnd1*, we simultaneously reduced *Lockd* using shRNAs (sh*Lockd*-1 and sh*Lockd*-2)(Figure S3A and S3B) and overexpressed *Ccnd1* with Ef1α-*Ccnd1* via *in utero* electroporation at E14.5, followed by the analysis of labeled cell subtypes at E16.5. Co-electroporation of sh*Lockd* and *Ccnd1*-overexpressed vectors was sufficient to rescue the reduction in the fraction of the PAX6^+^/TBR2^+^ cells to the control level and partially restored the increase in the PAX6^-^ /TBR2^-^ neuronal population in *Lockd*-KD (Figure 6A and 6B). Consistent with this, more *Lockd*-KD cells had differentiated into SATB2^+^ neurons and migrated into the cortical plate three days after IUE at E14.5 (Figure S5C and S5D). The introduction of *Ccnd1* overexpression significantly reduced the fraction of SATB2^+^ neurons in *Lockd*-KD and partially restored the SATB2^+^ neurons to the control level (Figure S5C and S5D).

**Figure 6.**
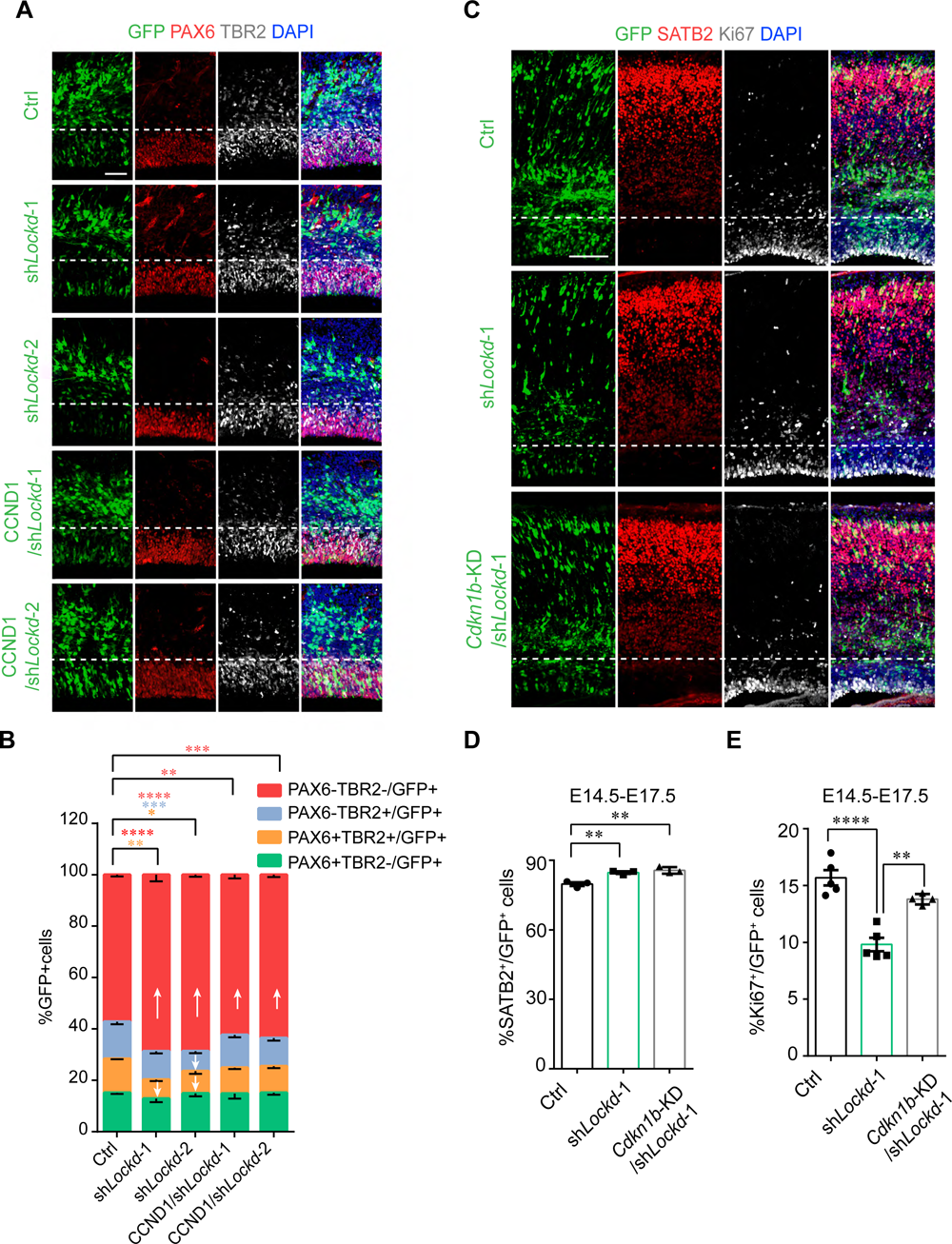
*Lockd* guarantees neural development through its *trans*-function on *Ccnd1* and *cis*-regulation for *Cdkn1b*. (A) Analysis of neural differentiation after single KD of *Lockd* or simultaneously with *Lockd* KD and *Ccnd1* overexpression. The indicated vectors (GFP^+^) were electroporated into the E14.5 cortex, and PAX6^+^ or TBR2^+^ cells were evaluated at E16.5 by IHC. Scale bar, 50μm. (B) Quantification of PAX6^+^ or TBR2^+^/GFP^+^ cell fractions in total GFP+ cells. The arrows indicate the up or down tendency relative to control group. Data represent mean ± SEM; statistical analysis is two-way ANOVA followed by Sidak’s multiple comparisons tests (n = 5; *p < 0.05, **p < 0.01, ***p < 0.001, ****p < 0.0001). (C-E) The shLockd (vector) or siRNA for knockdown of *Cdkn1b* (*Cdkn1b*-KD) were electroporated at E14.5, and the neuronal subtypes of the progeny cells or the precursor cells within the cell cycle (immunostaining Ki67 indicates proliferating precursor cells; SATB2 indicates UL neurons) were analyzed at E17.5. (C) Shown are the representative confocal images, the area below the dotted line is VZ-SVZ, the above area is IZ and CP. Scale bar, 100μm. The proportion of SATB2^+^ cells (D) (n=3) and the proportion of Ki67^+^ cells (E) (n=5, Ctrl; n=5, sh*Lockd*-1; n=4, *Cdkn1b*-KD/sh*Lockd*-1) in all the electroporated GFP^+^ cells were quantified. Values represent the mean ± SEM; statistical analysis is one-way ANOVA followed by Tukey’s multiple comparison test (**p < 0.01, ****p < 0.0001)

These results suggest that reduced expression of *Cdkn1b* in PlyA-KI can, to some extent, counteract the effect of down-regulated *Ccnd1*. We reasoned knocking down the RNA level of *Cdkn1b* with siRNA in *Lockd*-KD cells would relieve the phenotype to a similar level in PlyA-KI. Indeed, we found that knockdown of *Cdkn1b* in *Lockd*-KD restored the number of cycling NSPCs to the control level (Figure 6C, 6D, and 6E), indicating that *Cdkn1b* is necessary for pushing NSPC to exit the cell cycle. Notably, the degree of increased neuronal differentiation was not changed by *Cdkn1b*siRNA, suggesting that other factors could be involved in regulating neuronal differentiation.

## Discussion

The development of the cerebral cortex is an intricate but highly stereotyped process that is encoded by gene expression programs for the precise spatiotemporal control of neural stem/progenitor cells (NSPCs) self-renewal and differentiation. Although long non-coding RNAs (lncRNAs) have emerged as essential players in numerous physiological and pathological processes (Li and Chang, 2014; Wang et al., 2011; Wu et al., 2015), only a few lncRNAs have been genetic studied *in vivo*, especially in the context of neural development (Andersen et al., 2019; Li et al., 2019b). Here, we show for the first time that *Lockd* is an NSPC-enriched lncRNA, abundant in both the nucleus and cytoplasm. Furthermore, we combined PlyA-KI mediated premature termination of *Lockd* transcripts, and shRNAs mediated *Lockd* KD to explore the functions and genetic mechanisms of *Lockd* systematically *in vivo*. We found that *Lockd* functions *in cis* to regulate its upstream gene-*Cdkn1b* required for cell cycle exit and regulates *Ccnd1* essential for cell cycle progression *in trans* to guarantee the self-renewal and differentiation of NSPCs (Figure 7), offering the first example of lncRNA-mediated cell cycle control in NSPCs.

**Figure 7.**
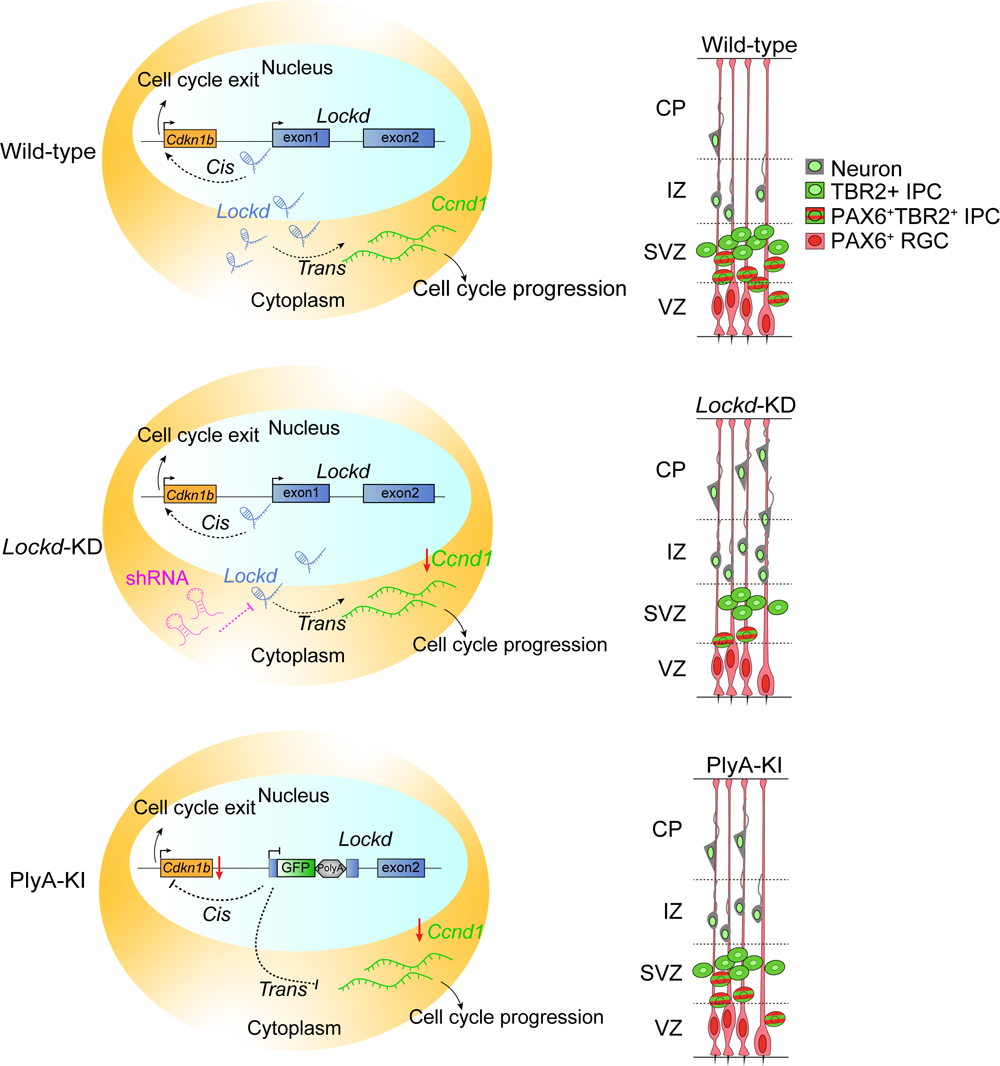
Model of *Lockd* function: The *cis*-and *trans*-actions of *Lockd* balance the proliferation and differentiation of NSPCs. In the WT, *Lockd* can regulate the cell cycle exit of NSPCs through its *cis* function on *Cdkn1b*; *Lockd* in the cytoplasm can regulate the abundance of *Ccnd1* via *trans* manner, thereby controlling the rate of cell cycle progression. The shRNA can delete the cytoplasmic transcripts of *Lockd*, which weaken its *trans* regulation on *Ccnd1* and promotes NSPC exit from the cell cycle to differentiate into neurons. However, PlyA-KI not only inhibits the *trans* regulation of *Lockd* but also disrupts its *cis*-regulation on *Cdkn1b*, thus, to some extent counteracting the effect of early withdrawal of cell cycle caused by repressed *trans* function. Ultimately, the phenotypes of PlyA-KI are less severe than that of *Lockd*-KD.

We initially characterized the expression pattern of *Lockd* by ISH on mouse brain sections and observed *Lockd* was highly expressed in the VZ-SVZ region. Further detection using SABER-FISH provided us a higher resolution for *Lockd* expression cell types: *Lockd* expressing cells also expressed *Pax6* and *Tbr2*. However, there was a more dynamic expression and specific spatial distribution for *Lockd*, as *Lockd* expression was much higher at the dorsal part of V-SVZ. Based on our findings and previous evidence of interkinetic nuclear migration (INM) of neural progenitors in the VZ, in which their nuclei migrate between the apical surface and the basal part of the VZ in concert with the cell cycle (Kosodo et al., 2011; Taverna and Huttner, 2010), we predict that *Lockd* is predominant during G1/G2 phase, and play a vital role in regulating cell cycle progression. Thus, identifying the *Lockd* expressing cell subtypes and their dynamic expression pattern will contribute to our understanding of molecular regulation of the cell cycle progression in NSPCs.

The biological significance of most lncRNAs during mammalian development is unclear due to the lack of genetic studies *in vivo* (Bassett et al., 2014; Nakagawa, 2016). To investigate the potential functions of *Lockd in vivo*, we chosen an EGFP-PolyA-tdTomato-PolyA cassette knock-in (PlyA-KI) strategy without deletion of the DNA sequence of *Lockd* locus. Similar genetic studies also use the insertion of PlyA signals into lncRNAs to block the transcriptional elongation and reveal *in vivo* phenotypes (Anderson et al., 2016; Bond et al., 2009). We found more PlyA-KI progenitors exit the cell cycle in advance, which led to reduced cycling progenitors and abnormal neuronal differentiation. Interestingly, the down-regulated genes in the PlyA-KI mouse cortex were preferentially associated with cell cycle progression, including *Cdkn1b*, suggesting a *cis* function of *Lockd*. Consistent with our findings, a recent study (Wang et al., 2019) assessed the lncRNA transcriptome in serum-starved mouse embryonic fibroblasts and established a bioinformatical model to predict the *Lockd*-targeting genes, which are involved in the processes of cell growth and cell proliferation.

However, *Cdkn1b* is not affected after PlyA insertion in the erythroid cell lines, where the *Lockd* promoter can physically associate with the *Cdkn1b* promoter, positively regulating *Cdkn1b* transcription through an enhancer-like *cis*-element (Paralkar et al., 2016). One possible reason is that the exon1 contains a regulatory DNA element in the cortex, and the insertion of a long polyA site-containing sequence can increase the distance between *Lockd* exon1 and *Cdkn1b* promoter, thereby potentially altering their activity. Nevertheless, the histone modifications in *Lockd* exon1 indicated a promoter-like element, and we were unable to find any enhancer-like element within the *Lockd* exon1. Although PlyA-KI of *Lockd* should not impair its possible DNA-element dependent *cis* functions, the strategy can disrupt transcriptional elongation, and such transcriptional activity nearby the TSS region can also regulate neighboring genes in an enhancer RNA (eRNA) mechanism (Elling et al., 2018). The enhancer RNA (eRNA) originates from transcriptionally active enhancer regions (H3K4me1^high^ H3K4me3^low^ H3K27ac^high^) and functions *in cis* to enhance the expression of neighboring protein-coding genes (Creyghton et al., 2010; Heintzman et al., 2007; Visel et al., 2009). We propose that the transcriptional activity local to the TSS of *Lockd* is essential, and *Lockd* can regulate upstream gene-*Cdkn1b* through a non-classical eRNA mechanism because *Lockd* exon1 is produced from an active promoter region (H3K4me3^high^) instead of active enhancer regions. Similar to *Lockd*, the *lincRNA-Cox2* (long intergenic non-coding RNAs) locus has both high H3K4me1 and H3K4me3 marks, distinct from the typical definition of eRNAs, also controls the expression of the neighboring gene *Ptgs2 (Cox2) in cis*(Elling et al., 2018). Future work aimed at knocking down the expression of *Lockd* using CRISPRi, which mediates heterochromatin formation and silencing of gene transcription, could confirm the importance of *Lockd* transcription for *Cdkn1b* levels.

Deletion of *Cdkn1b* in mouse (*Cdkn1b*^-/-^) leads to enhanced growth, multiple organ hyperplasia, retinal dysplasia, and pituitary tumors (Kiyokawa et al., 1996; Nakayama et al., 1996), and overexpression of *Cdkn1b* promotes the cell cycle exit of NSPCs (Tarui et al., 2005). However, PlyA-KI mice exhibited a deficit in cell cycle progression, especially for the advanced cell cycle exit, which is different from *Cdkn1b*^-/-^. As PlyA-KI blocked both the transcription of the *Lockd*exon1 locus and the production of mature lncRNA *Lockd*, making it difficult to distinguish between the *cis* and potential *trans* functions for this gene. *Lockd* is abundant in both the nucleus and cytoplasm. Therefore, we use shRNAs to efficiently knockdown (KD) the *Lockd* levels in the cytoplasm to explore its potential *trans* functions. The *Lockd*-KD in the cortex showed dramatically decreased transitioning IPs and increased neuronal differentiation. Further RNA-seq analysis comparing PlyA-KI with *Lockd*-KD showed an overlapped DEGs regulating cell cycle progression and neural differentiation. Not surprisingly, we also observed a broad set of non-overlapped DEGs, such as *Cdkn1b,* only downregulated in PlyA-KI, and we believe that this is since *Lockd*-KD reveals only the functions of *Lockd* in *trans*, whereas the PlyA-KI mice exhibit both *cis* effects on *Cdkn1b* and *trans* effects on other cell cycle genes. Furthermore, several studies show that shRNA can also modulate local gene expression (Luo et al., 2016) (Orom et al., 2010; Wang et al., 2011); we can not exclude the possibility of transcripts exchange between the nucleus and cytoplasm after *Lockd*-KD. Therefore, we should integrate multiple approaches to investigate the function and molecular mechanism of *Lockd* in the future. For example, antisense oligonucleotides (ASOs) can target RNAs in the nucleus and cytoplasm, disturbing both *cis* and *trans* functions of lncRNAs (Liang et al., 2011). Overexpression of lncRNAs by IUE or generation of transgenic mice that express lncRNAs at physiological levels from an integrated bacterial artificial chromosome (BAC) construct (Andersen et al., 2019) can indicate the *trans* functions of lncRNAs.

Previous evidence suggests that the cell cycle regulation, specifically of the G1 phase, plays a crucial role in controlling the self-renewal versus neuronal differentiation of neural progenitors (Dehay and Kennedy, 2007; Lange et al., 2009). From our RNA-seq data, *Cyclin D1* (*Ccnd1*) were both downregulated in PlyA-KI and *Lockd*-KD mice, and *Ccnd1^-/-^* mice display developmental abnormalities, hypoplastic retinas, and pregnancy-insensitive mammary glands (Fantl et al., 1995; Sicinski et al., 1995), which contrasts with the phenotypes in *Cdkn1b^-/-^*, but is in line with our *Lockd*-KD phenotypes of reduced proliferation and increased neuronal differentiation. Consistent with this, overexpression of *Ccnd1* can rescue the defects of proliferation in *Lockd*-KD, suggesting *Ccnd1* as the downstream target regulated by *Lockd in trans*. Given the opposite phenotypes between *Ccnd1^-/-^* and *Cdkn1b^-/-^*, crossing *Ccnd1^-/-^* with *Cdkn1b^-/-^* mice can restore the developmental defects of *Ccnd1^-/-^* mice to wild-type phenotypes (Geng et al., 2001; Tong and Pollard, 2001). Both *Ccnd1*and *Cdkn1b* were downregulated in PlyA-KI mice, but they still showed defects in the cell cycle of NSPC, although weaker than *Lockd*-KD. A recently published study found that lentiviral or siRNA-mediated knockdown of *Lockd* transcripts in NIH3T3 cell lines and primary fibroblasts resulted in significant downregulation of a group of kinesin superfamilies (Kif4, Kif11, and Kif14) known to be involved in mitosis (Johnsson et al., 2020), which is consistent with our *Lockd*-KD sequencing results (Figure 5D). Notably, CDKN1B binds to the Kif11 promoter through a p130-E2F4-dependent mechanism as a transcriptional repressor of Kif11 (Pippa et al., 2012). Kif14 affects the proteasomal degradation pathway of CDKN1B by regulating the ubiquitination level of CDKN1B (Xu et al., 2014). Therefore, we propose a hypothesis that *Lockd*-KD does not inhibit *Cdkn1b* transcription but reduces Kif14 expression, thereby hindering the degradation process of CDKN1B. The increased level of CDKN1B protein further inhibits *Kif11* expression, which then promotes cell cycle exit. PlyA-KI model blocks both *cis* and *trans* functions of *Lockd*, causing enhanced stability of CDKN1B protein by the repressed *trans* function and reduced *Cdkn1b* transcription due to the inhibited *cis* role. Such a combined effect lead to a less severe phenotype in PlyA-KI than that in the *Lockd*-KD model. PlyA-KI mice were viable, healthy, and fertile, indicating the transient and spatiotemporal function of *Lockd* without influencing the overall fitness of animals. Similar to *Lockd*, *Evx1as* is transiently required in the patterning of the primitive streak but did not lead to a substantial effect on embryonic development (Han et al., 2018).

Because lncRNAs can function *in cis* and *trans* to regulate the same or different genes, making it much more complex and challenging to investigate the potential mechanism than protein-coding genes or microRNAs (Liu and Lim, 2018). The lncRNA *Haunt* acts to prevent aberrant *HOXA* expression, and its genomic locus contains potential enhancers of *HOXA* activation (Yin et al., 2015). Importantly, *cis* and *trans* mechanisms for lncRNAs are not necessarily mutually exclusive, and some lncRNAs can also carry out multiple distinct molecular roles through both manners (Cajigas et al., 2018; Pavlaki et al., 2018; Vance et al., 2014). Future work to comprehensively identify proteins that interact with *Lockd* will be necessary to determine the mechanisms and relationships between *cis* and *trans* functions, by which this lncRNA regulates neural development.

Together, our work contributes new data for a better understanding of how lncRNAs can function both *in cis* and *in trans* to equilibrate NSPCs expansion and differentiation, which is crucial for brain development, perhaps the regulation of cortical expansion during development and evolution (Fish et al., 2008; Kriegstein et al., 2006). Although due to the positive and negative selection during evolution, most lncRNAs are not conserved in their primary sequence or even only detected in specific species (Briggs et al., 2015), some experimental studies have now shown that the function of specific lncRNAs can be preserved regardless of the sequence conservation (Pang et al., 2006; Ulitsky et al., 2011). There are thousands of different lncRNAs expressed in the developing brain, and emerging studies have shown the relevance between lncRNAs and neural disease pathogenesis (Briggs et al., 2015; Meng et al., 2015). Therefore, combinational studies of multiple genetic animal models with gene-regulating tools (RNAi and overexpression) to dissect how *cis* and *trans* functions of lncRNAs regulate brain development and contribute to neurological disorders is an excellent area for future research.

## STAR⁎METHODS

### EXPERIMENTAL MODEL AND SUBJECT DETAILS

#### Maintenance of mice

All mice were group-housed and maintained according to the Tongji University Guide for the use of laboratory animals. The mouse was housed under specific pathogen-free conditions at 22 ± 2°C and fed with free access to standard mouse chow and tap water. The noon of the day on which the vaginal plug is found is counted as embryonic (E) day 0.5.

#### *Lockd* PlyA-KI mice

*Lockd* PlyA-KI mice were generated by Cyagen Bioscience Incorporated Company (Suzhou, China). Briefly, the gRNA targeting the 80 bp downstream of *Lockd* TSS, the donor vector containing the loxP–EGFP–WPRE–polyA–loxP–tdTomato–WPRE-polyA cassette, and Cas9 mRNA were co-injected into fertilized C57BL/6 background mouse eggs to generate targeted conditional knockin offspring (Figure 2B). F0 founder animals were identified by PCR followed by sequence analysis, which was bred to wildtype mice to test germline transmission and F1 animal generation.

#### Cell lines

Neuro-2a (N2a) cells purchased from Peking Union Medical College Hospital and HEK293T cells (Kee. K lab, Tsinghua University) were maintained in indicated culture media (MEM or DMEM) containing 10% fetal bovine serum (Thermo Fisher Scientific) and 1% penicillin/streptomycin (Life Technologies).

## METHOD DETAILS

### DNA constructs and siRNA

For knockdown experiments, shRNA sequences (see Key Resources Table) were cloned into the FUGW-H1 lentiviral plasmid expressing GFP or TurboRFP under the control of the human ubiquitin C (hUbc) promoter and the specific shRNA driven by human U6 promoter, and lentiviruses were generated as described previously (Hu et al., 2017). The efficacy of each shRNA was confirmed in Neuro-2a cell lines (Peking Union Medical College Hospital) infected by lentivirus for 48h before RNA extraction. For constructing *Lockd* overexpressing vectors, full-length mouse *Lockd* (Paralkar et al., 2016) was amplified by polymerase chain reaction (PCR) from cDNA of the E13.5 cortex, then subcloned into the pCAG vector expressing *Lockd* under the control of CAG promoter followed with a bovine growth hormone polyadenylation signal and EGFP under the control of hUbc promoter. For constructs of *Ccnd1*, the coding sequence for *Ccnd1* was amplified by PCR from cDNA of the E13.5 cortex and was cloned into the pEF1α -IRES-Zsgreen in frame with sequence encoding N-terminal 3× Flag tag. Primer information has been included in the Key Resources Table. The p*NeuroD1*-GFP plasmid was a gift from Dr. Frank Polleux (Guerrier et al., 2009).

The following siRNAs were purchased from Ambion: 5’ - GCUUGCCCGAGUUCU ACUAtt - 3’ (sense strand) and 5’-UAGUAGAACUCGGGCAAGCtg -3’ (antisense strand) for *Cdkn1b* siRNA, predesigned (no. 118712) and verified *in vivo* (Nguyen et al., 2006); The control siRNA (Silencer Negative Control# 1 siRNA no. 4611) with no significant hom ology with known gene sequences in the rodent.

### In situ hybridization (ISH)

Protocols for ISH were detailed previously (Liu et al., 2019). Briefly, the RNA probe sequence was PCR amplified from cDNA of the E13.5 cortex and cloned into the pBluescript II KS vector (a gift from Dr. Zhengang Yang, Fudan University) or pEASY-T3 cloning vector (TransGen, CT301-01). Primers used were included in the Key Resources Table. Digoxigenin-labeled riboprobes were transcribed using the DIG-RNA Labeling Mix (Roche). Embryonic heads (E11.5–E13.5) or isolated brains (E14.5 and older) were fixed at 4°C in 4% PFA for 4-5 hr or overnight depending on the stage as previously described (Hu et al., 2017). Brains were equilibrated in 30% sucrose/PBS and frozen in Tissue Tek OCT (Sakura Finetek, Torrance, CA). Cryosections (20 μm) were incubated with the DIG-labeled RNA antisense probe (0.1ng/μL in hybridization buffer) overnight at 65 °C, washed in 0.2× SSC three times for 20 min each at 65 °C and once at room temperature (R/T). Sections were blocked with 10% sheep serum (Sigma) in buffer B1(0.1 M Tris-HCl, pH 7.4; 0.15M NaCl) for 1 hour at R/T and then incubated with anti-DIG antibody (Roche; 1:5000) overnight. Sections were washed with buffer B1 four times for 20 min each, prestained twice with buffer B2 (0.1M Tris-HCl, pH 9.5; 0.1M NaCl; 0.05M MgCl2; 0.1% Tween-20) for 5 minutes each time at R/T, followed by staining with 50x NBT/BCIP (Roche) in buffer B2 for 2-4 hr. The reaction was stopped and washed in PBS three times for 15 min and mounted with fluoro-mount GTM (Southernbiotech).

### Fluorescent RNA in situ hybridization (FISH)

FISH probes were designed and applied using the original protocol (Kishi et al., 2019) based on signal amplification by exchange reaction (SABER). In brief, the BED files for *Pax6* (NM_001244200.2), *Tbr2* (NM_010136.3), and *Lockd* (NR_037955.1) were generated through the UCSC Table Browser. Genome-wide probe set for the mouse was downloaded from The Wu lab Oligopaints website (https://oligopaints.hms.harvard.edu/) with a Balance setting. The overlapping probes between the BED output file and the genome-wide chromosome-specific BED file were identified with intersectBed (bedtools 2.27.1). The oligos were ordered from Sangon Biotech and pooled together at a final amount of 10nmol, resuspended in distilled Water to 10 µM.

A Primer Exchange Reaction concatemerization reaction (1X PBS; 10 mM MgSO_4_; dNTP, 0.6 mM of A, C, and T, NEB-N0446S; 0.1 µM Clean.G; 800 U/ml Bst Polymerase, NEB-M0374L; 0.5 µM hairpin; 1 µM oligo pool) was applied to extend the pooled oligos. Reactions were incubated as follows: 15 min at 37°C without oligo pool; adding oligo pool for an extension of 2 hr at 37°C; 20 min at 80°C; 4°C holds. The reaction products were purified and concentrated using a MinElute kit (Qiagen, cat. no. 28004) with distilled-water elution and then confirmed on a 1% agarose gel with ethidium bromide. The probe sets of 300–700 nt were used for the study. Fluorescent oligos were ordered from Sangon Biotech with a 5’ modification of Cy3 for *Lockd* detection and Quasar 670 for *Pax6* or *Tbr2* detection.

For SABER-FISH on the mouse cortex, E15.5 mouse brains were dissected and fresh-frozen with OCT in a liquid nitrogen bath. Cryosections of the brain were prepared at 13 µm thickness and permeabilized in PBS/0.5% Triton X-100 for 20 min at R/T. The sections were subsequently fixed in 4% PFA for 20 minutes and washed twice in PBS for 5 min each at R/T. Then, the PBS was replaced with pre-warmed (43°C) 40% wHyb (2x SSC, Invitrogen; 40% deionized formamide, Invitrogen; 1% Tween-20, Sangon; diluted in UltraPure Water) for at least 15 min at 43°C. Samples were then incubated with 120 µL of pre-warmed (43°C) probe mix (1 µg of probe per gene; 96 µL of Hyb1 solution (2.5x SSC; 50% deionized formamide; 12.5% Dextran Sulfate, Ameresco; 1.25% Tween-20); diluted up to 120 µL with UltraPure Water) and incubated 16–48 hr at 43°C. The sections were washed twice with 40% wHyb at 43°C for 30 min each, twice with 2x SSCT (2x SSC/ 0.1% Tween-20) for 10 min each. For branching (used to amplify signals of *Lockd*) after primary probe washes, the branch probe was extended to a length of 500 nt and incubated for 12-16 hr in the Hyb1 solution at 37 °C. Washes were performed as for the primary probe incubation with the temperature set to 37 °C. For fluorescent detection, slides were washed twice in PBSTw (D-PBS, Sangon; 0.1% Tween-20) at 37 °C, and then were incubated with 120 µL of the fluorescent oligo mix (0.2 μM fluorescent oligo diluted in a 1× PBS solution with 0.2% Tween-20 and 10% dextran sulfate.) for 2 hr at 37°C. The samples were subsequently washed in PBSTw three times at 37°C and counterstained with DAPI. Slides were mounted with anti-fade fluoro-mount GTM. A full list of oligo sequences for every gene-specific probe set and sequences for fluorescent oligos, primers, hairpins, and branches used in this study were listed in Table S1.

### Subcellular fractionation

Subcellular fractionation was performed as previously described (Cabianca et al., 2012). E13.5 mouse cortices were dissected, dissociated, and then collected by centrifuging at 400 x g for 10 min (Qian et al., 2000; Shen et al., 2004). The pellet was lysed with 200 μl/10^6^ cells of cold RLN1 solution (50 mM Tris HCl pH 8.0; 140 mM NaCl; 1.5 mM MgCl2; 0,5% NP-40; 2mM Vanadyl Ribonucleoside Complex, Sangon Biotech; reagent stocks were prepared in RNase-free H20) for 5 min on ice. For N2a, 2.5 x 10^5^ cells were used. After centrifuging at 4 °C, 300 ×g for 2 min, the supernatant was transferred into a new tube, storing as the cytoplasmic fraction. The remaining pellet corresponds to the nuclear fractions.

### Gene expression analysis

Total RNA from whole cell lysate or specific subcellular fractions were extracted using TRIzol reagent (Invitrogen) according to the manufacturer’s instructions. Equal amounts of RNA (100 ng-1 μg) were reverse-transcribed into cDNA using TransScript One-Step gDNA Removal and cDNA Synthesis SuperMix (TransGen Biotech, AT311). Quantitative real-time-PCR (qRT-PCR) was carried out in triplicates on a Bio-Rad CFX384 Touch Thermal Cycler using ChamQTM SYBR qPCR Master Mix (Vazyme) following the manufacturer’s instructions. The data were calculated using the delta-delta C.T. method with *Gapdh* as a control gene or presented as a percentage of the total amount of detected transcripts. The primers used in the qRT-PCR are shown in Table S1.

### In utero electroporation (IUE)

In utero microinjection and electroporation were performed as described (Hu et al., 2017). In brief, timed-pregnant ICR mice (Vital River Laboratory) at E14.5 were anesthetized, and the uterine horns were exposed. Approximately 0.5-1 μL of plasmid solution (2-3 μg/μl) was injected manually into the lateral ventricle of the embryonic brain using a micropipette. Five pulses (36 V, 50 ms in duration with a 999 ms interval) were delivered across the uterus with a homemade 10 mm-diameter disc electrodes by a square wave electroporator (CUY21VIVO-SQ; BEX). After electroporation, uterine horns were then placed back into the abdominal cavity, and the wound was sutured to allow the embryos to continue development. Mouse embryos were analyzed at E16.5 or E17.5. In rescue experiments of *Lockd* knockdown, sh*Lockd*-1,2 and *Ccnd1* overexpression plasmids were mixed (2.5 μg/μl each) and injected into the lateral ventricle. The same ratio used in other co-electroporation experiments included sh*Lockd*-1 and p*Neurod1*-EGFP. For RNAi experiments, siRNAs were injected at 20 μM concentration together with GFP expression plasmids (shCtrl, sh*Lockd*-1) at 2 μg/μl concentration.

### Bromodeoxyuridine (BrdU) labeling

For BrdU (5-Bromo-2’-deoxyuridine, Sigma) labeling, timed pregnant mice were administrated with BrdU (100 μg per gram of body weight) by intraperitoneal injection at defined time points before sacrifice.

### Measurement of cell cycle kinetic parameters

The total cell cycle duration (Tc) and the length of S-phase (Ts) of the NSPC in the VZ were determined by BrdU/EdU dual pulse labeling method adapted from Martynoga et al. (Martynoga et al., 2005)and Harris et al. (Harris et al., 2016). In brief, pregnant mice (E14.5) were injected intraperitoneally with EdU (50 mg/kg) at T = 0 hr to label all S-phase cells at the beginning of the experiment. 1.5 hr after this (interval duration Ti), another injection of BrdU (50 mg/kg) was given. The animal was sacrificed 0.5 hr after BrdU injection and embryos immersion-fixed in 4% PFA. Sections were stained for BrdU (B35128, ThermoFisher), and the detection of EdU was performed with Click-iT assay (Invitrogen) using Alexa Fluor 647 azide according to the manufacturer’s instructions. As the length of the cell cycle duration is directly proportional to the number of cells in their corresponding period (Nowakowski et al., 1989); therefore, the Ts can be calculated as Ts = Ti * (EdU^+^BrdU^+^ cells/EdU^+^BrdU^−^ cells), and Tc = Ts / (EdU^+^BrdU^+^ cells/total VZ progenitors).

### Analysis of cell cycle progression by EdU pulse labeling

Analyses of cell cycle progression of mouse NPCs were performed based on a previously described protocol (Wang et al., 2014; Yoon et al., 2017). Briefly, mouse NPCs from E13.5 WT and PlyA-KI forebrain were dissociated and cultured in serum-free N-2/B-27/DMEM medium with 10 ng/ml FGF2 (Life Tech) in poly-L-lysine (PLL, Sigma)-coated 6 well culture plates (Costar) for 2 days. Then, NPCs were pulsed by 10 μM EdU (ThermoFisher) for 30 min and washed thoroughly with NPC media. After culturing for 5 hr, cells were dissociated by Accutase, fixed with 4% paraformaldehyde in PBS for 20 min at 4°C, stained with Click-iT EdU Alexa 647 Flow Cytometry Kits (ThermoFisher) according to the manufacturer’s instructions, and stained with FxCycle™ Violet (ThermoFisher) for DNA content. The cells were then applied to flow cytometry using CytoFLEX (Beckman). Approximately 8000-10000 cells were analyzed for each group (WT and PlyA-KI) in every biological replicate.

### Immunohistology and confocal imaging

Brain section: following dissection, brains were fixed at 4°C with 4% paraformaldehyde (PFA) in PBS as previously described (Hu et al., 2017), and then were transferred to 30% sucrose/PBS solution at 4 °C for cryoprotection. Brains were embedded and frozen in the OCT and sectioned coronally (20 μm-thickness) on a Leica CM1950 cryostat. For staining, cryostat sections were rehydrated with PBS, blocked, and permeabilized in 5% BSA/PBST (1 x PBS, 0.3% Triton X-100) for 1 hr at room temperature, followed by incubation with primary antibodies at 4°C overnight. After washing with PBS, fluorescence-conjugated secondary antibodies were applied to the sections for 2 hr at R/T. After rinse, 0.1 mg/ml 4,6-diamidino-2-phenylindole (DAPI, Thermo Fisher Scientific) in PBS was added for nuclei staining. For BrdU labeling, sections were treated with 20 μg/mL proteinase K (Sigma) for 5 min, followed by 250 U/ml DNase Ⅰ (Sangon Biotech) for 10 min at R/T before BrdU staining. Stained sections were mounted with anti-fade fluoro-mount GTM. All the antibodies used are listed in the Key Resources Table.

### Fluorescent-activated cell sorting (FACS)

Two days after electroporation, GFP transfected cortices from 3–5 embryos of E16.5 were microdissected and dissociated as previously described (Hu et al., 2017; Liu et al., 2019). GFP-positive cells were isolated using the FACS Aria Ⅱ (BD Biosciences) system. Gating for GFP positive cells was set using nontransfected cortical cells. Cells were directly sorted into hibernation medium (30mM KCl, 5mM NaOH, 5mM NaH_2_PO_4_, 0.5mM MgCl_2_ꞏ6H2O, 20mM Sodium Pyruvate, 5.5mM Glucose, 200mM Sorbitol, pH7.3-7.4, filtered with 0.22 μm filter).

### RNA-sequencing and bioinformatic analyses

Following total RNA isolation from FAC-sorted GFP-positive cells using TRIzol reagent (Invitrogen), RNA was quality controlled and quantified using an Agilent 2100 Bioanalyzer. Next, the sequence libraries were generated using the KAPA Hyper Prep Kit for the Illumina platform (kk8504), following the manufacturer’s instructions. Paired-end 150-bp (PE150) sequencing was further performed on a HiSeq X10 (Illumina) at Berry Genomics. For transcriptome profiling of mouse developing forebrains, cortex from W.T. or PlyA-KI E13.5 embryos were dissected and followed by RNA extraction as described above. The Annoroad Corporation performed the strand-specific sequencing libraries preparation and PE150 sequencing on the NovaSeq 6000 (Illumina) platform. The generated sequences of clean data were mapped against the mouse reference genome (GRCm38/mm10) using HISAT2 v2.1.0 (Kim et al., 2015). Read counts generated by HISAT2 were analyzed for differential expression via DESeq2 (Love et al., 2014) (version 1.16.1). The genes with both fold change greater than 1.2 and P-values less than 0.05 between two groups were identified as significant differential expressed genes (DEGs). Cufflinks (Trapnell et al., 2010) were used to calculate the relative transcript abundances (FPKM, Fragments Per Kilobase of transcript per Million mapped reads). Functional annotation and gene ontology analyses of DEGs were based on the Metascape database (Zhou et al., 2019). Circos plot was created using ClicO FS (Cheong et al., 2015).

## QUANTIFICATION AND STATISTICAL ANALYSIS

### Imaging analysis and cell counting

Imaging of the GFP signal on the embryonic PlyA-KI mouse brain section was conducted on an inverted Zeiss Axio Observer Z1 with an LED light source (Figure 2C). If not explicitly indicated, remaining tissue images were acquired using a Zeiss LSM 880 microscope, and SABER-FISH was imaged with the AiryScan detector. Maximum-intensity projections in Z were created using ZEN software from raw multichannel Z-stacks.

For the quantitative analysis of IUE data, the electroporated region of each coronal section was pictured and compared with the equivalent section in the littermate counterparts. Only electroporated cells within the dorsolateral cortex were examined. The marker-positive cells were defined and counted based on the intensity of marker immunofluorescence in electroporated cells measured with the same criteria among different groups, and the percentage of marker positive cells in electroporated cells was calculated. For quantification of cell cycle progression in WT and PlyA-KI mice at E17.5, the anatomically matched regions were identified, and only BrdU+ cells within each vertical column of 200 μm width and 125 μm distance from the ventricular surface were measured. For distribution plots of FISH puncta, puncta within 200 μm distance from the ventricular surface were measured, and the numbers of FISH puncta per 200 x 125 μm^2^ area in every 20 μm interval from the ventricular surface was plotted as a histogram of percentage. Quantifications were performed using Imaris software (Bitplane) or ZEN software (blue edition, Zeiss). All quantifications were performed with 4-10 brain sections from at least 4 animals. The data in this study are presented as the means ± SEM, and the statistical significance was determined using Student’s t test, one-way or two way ANOVA followed by Dunnett’s post-test or Sidak’s multiple comparisons test accordingly and reported as *p < 0.05, **p < 0.01, ***p < 0.001, ****p < 0.0001 (GraphPad Prism software). The statistical details of each experiment can be found in the relevant figure legends.

## DATA AND SOFTWARE AVAILABILITY

All software used in this study is freely or commercially available (see the Key Resources Table). The accession number for the RNA-seq datasets reported in this paper is NCBI GEO: GSE∼∼∼.

## ACKNOWLEDGMENTS

We thank Dr. Frank Polleux for providing p*NeuroD1*-GFP plasmid; Dr. Zhengang Yang for pBluescript II KS vector and help in the experiment of *in situ* hybridization. Drs. Peng Yin, Jocelyn Y. Kishi, Brian J. Beliveau for technical assistance in SABER-FISH assay; Dr. Jia Li for providing RNA-seq data of Ebf2-EGFP E11.5 mouse forebrain; Huan Wu, Dr. Qing-Ran Bai for their help in several experiments; members of the Shen laboratory for suggestions and comments. The research was supported by the medical school of Tsinghua University and the school of life science and technology of Tongji University.

## AUTHOR CONTRIBUTIONS

S-J.Q., J.Z. designed and conducted experiments and performed data analysis; Q.-R.B. helped with plasmids construction; H.W. helped with experiments and breeding of PlyA-KI mouse lines. Q.S. supervised the project. S-J.Q. and Q.S. wrote the manuscript.

## Supplemental Information

**Figure S1.**
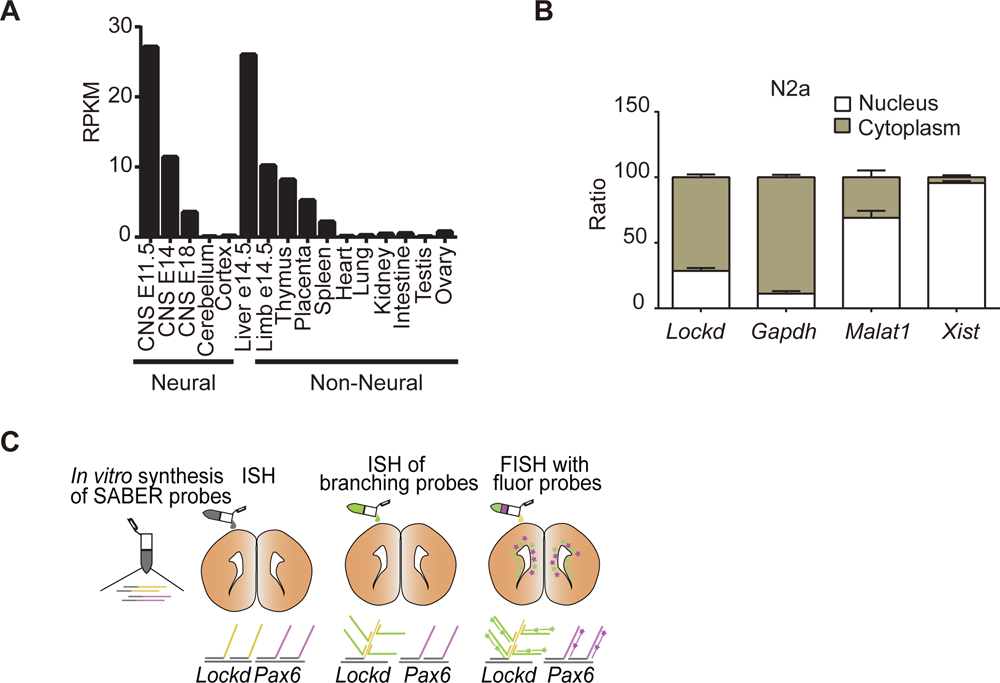
Characterization of *Lockd*, a lncRNA Enriched in NSPCs, Related to Figure 1. (A) *Lockd* prominently expresses in neural tissues and some non-neural organisms like the liver and thymus. Data collect from NCBI. Reads per kilobase per million mapped reads (RPKM). (B) Subcellular fractionation of indicated lncRNAs and mRNAs in N2a cell lines followed by RT-qPCR. Error bars depict mean ± SEM from three biological replicates. (C) Workflow for SABER-FISH detection of *Lockd* and *Pax6* transcripts. First, probes targeting *Lockd* and *Pax6* with different concatemer sequences on their 3’ ends are synthesized in *vitro*. These probes are then hybridized to *Lockd* and *Pax6* simultaneously in E15.5 mouse brain sections. For *Lockd* transcripts, another round of branching probes binding can be used to increase the signal further. After that, fluorescently labeled oligonucleotides can be hybridized to complementary concatemers.

**Figure S2.**
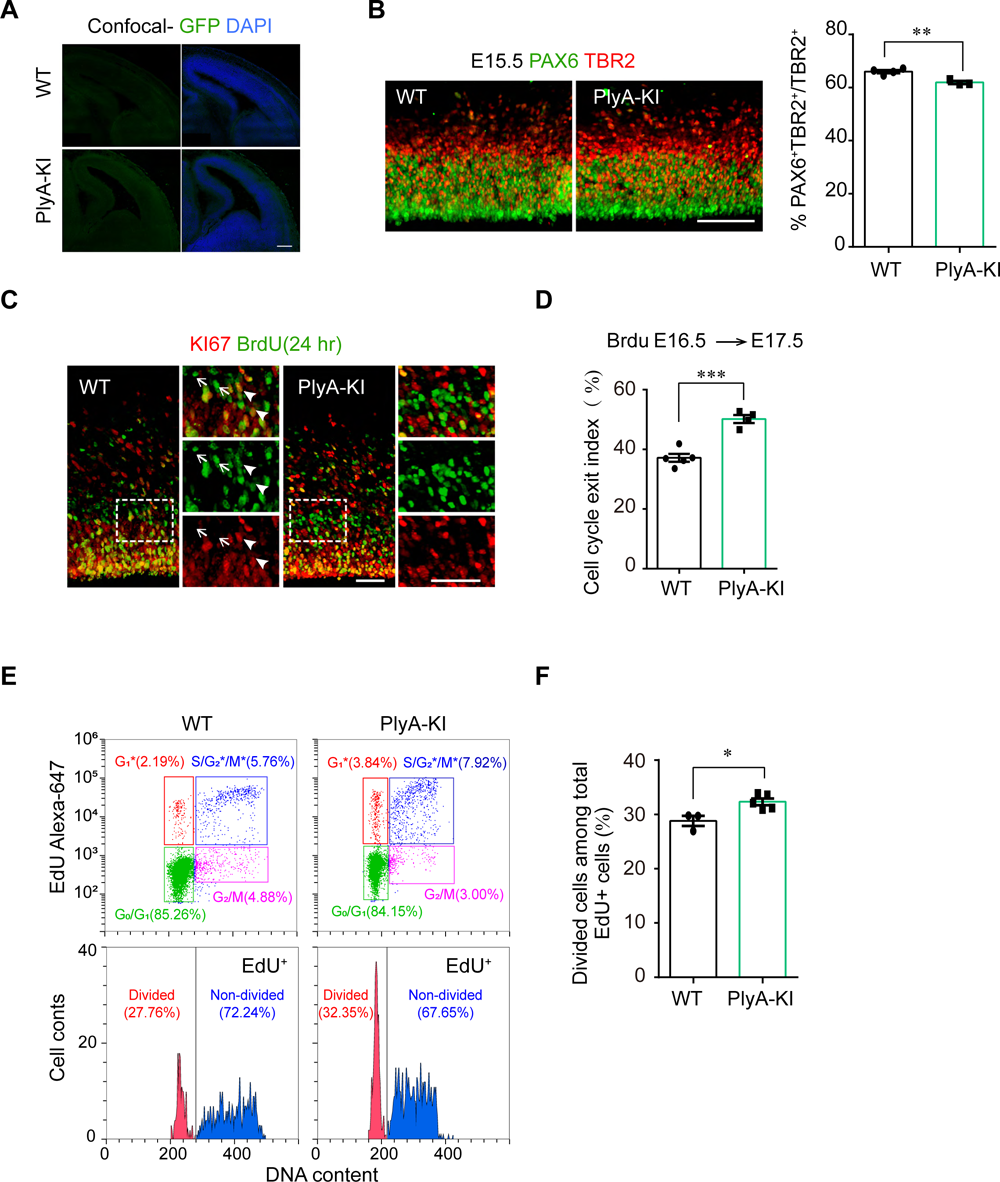
Generation of a mouse line with *Lockd* premature termination and analysis of *Lockd* functions *in vivo,* Related to Figure 2. (A) Represent images of the GFP immunostaining in E13.5 WT and PlyA-KI cortex pictured by LSM-880 confocal microscopy. Scale bar, 200 μm. (B) E15.5 WT and PlyA-KI cortical sections immunostained for PAX6, TBR2 (left, scale bar, 100μm). The right plot shows the fraction of PAX6^+^/TBR2^+^ cells relative to total TBR2^+^ cells. Values represent mean ± SEM (n = 4, WT; n = 3, PlyA-KI; **p < 0.01; unpaired Student’s t test). (C and D) Cell cycle exit analysis. Pregnant mice were injected with BrdU at E16.5 and analyzed 24 hrs later. Arrowheads point to Ki67^+^/BrdU^+^ cells, and arrows indicate examples of Ki67^−^/BrdU^+^ cells (C, scale bar, 50 μm). The fraction of KI67^−^/BrdU^+^ cells relative to BrdU^+^ cells was quantified (D). Values represent mean ± SEM (n = 5, WT; n = 4, PlyA-KI; ***p < 0.001; unpaired Student’s t test). (E and F) Flow cytometry analysis of cell cycle progression of WT and PlyA-KI NPCs. Mouse NPCs were cultured from the E13.5 cortex for 2 days and pulse-labeled with EdU (10 μM) for 30 mins; After 5 hrs incubation, EdU and DNA content (FxCycle™ Violet) were stained, followed by flow cytometry analysis. Upper panels of (E) are representative sample dot plots at 5 hrs after EdU labeling. Color boxes indicate the cells in a specific cell cycle phase. The EdU^+^ cells were grouped into divided (G1*) and non-divided (S/G2*/M*) populations. The sample histograms of DNA content from EdU cells are shown in the lower panels of (E) and quantification (F). Values represent mean ± SEM (n = 3, WT; n = 5, PlyA-KI; *p < 0.05; unpaired Student’s t test).

**Figure S3.**
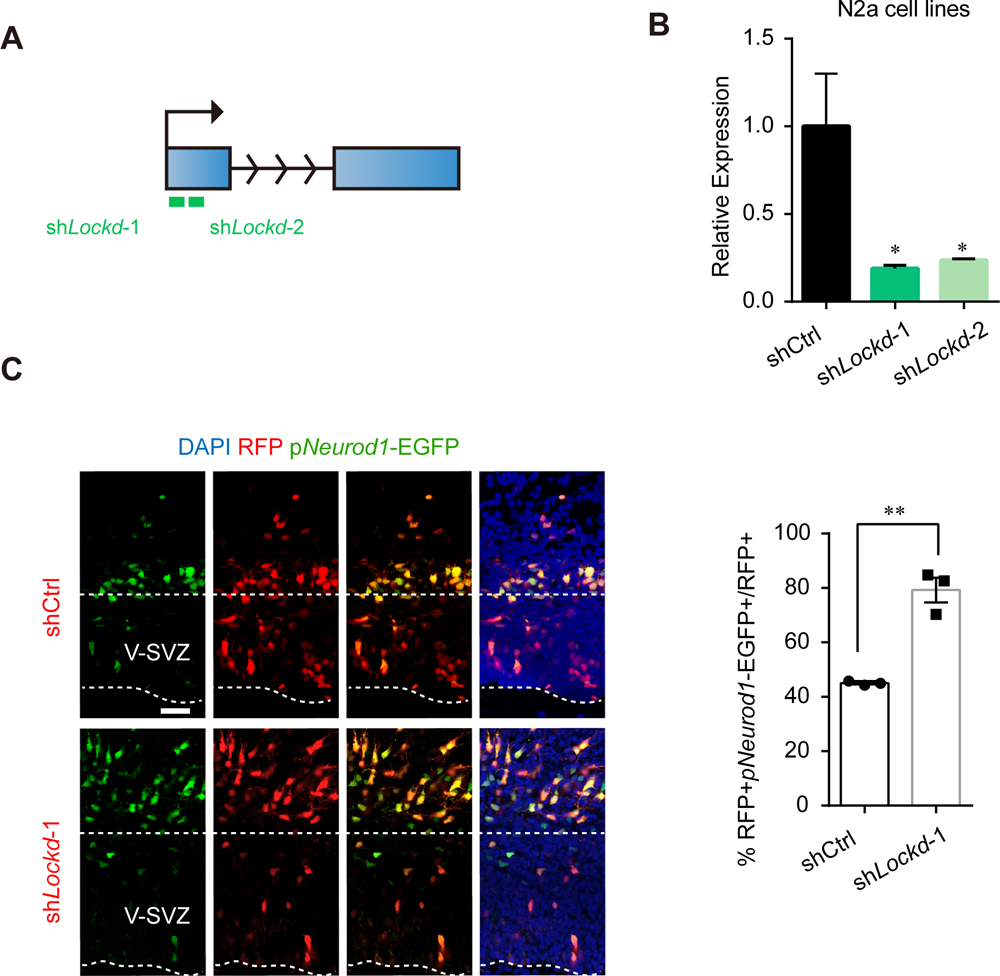
Identify the *Lockd* knockdown phenotypes, Related to Figure 4. (A) Schematic diagram of *Lockd* RNA and the regions targeted by sh*Lockd*-1 and sh*Lockd*-2. (B) RT-qPCR analysis of two distinct shRNAs targeting *Lockd*(sh*Lockd*-1 and -2) in Neuro2a cell lines. Values represent mean ± SEM (n=3; *p < 0.05; one-way ANOVA followed by Dunnett’s multiple comparisons test). (C) Represent images of cortex sections electroporated at E14.5 with shCtrl-RFP or sh*Lockd*-RFP and p*Neurod1*-EGFP constructs. Scale bar, 50μm. Right: quantification of RFP^+^, EGFP^+^ cells as a percentage of total RFP^+^ cells. Data represent represent mean ± SEM (n=3; **p < 0.01; Unpaired Student’s t test).

**Figure S4.**
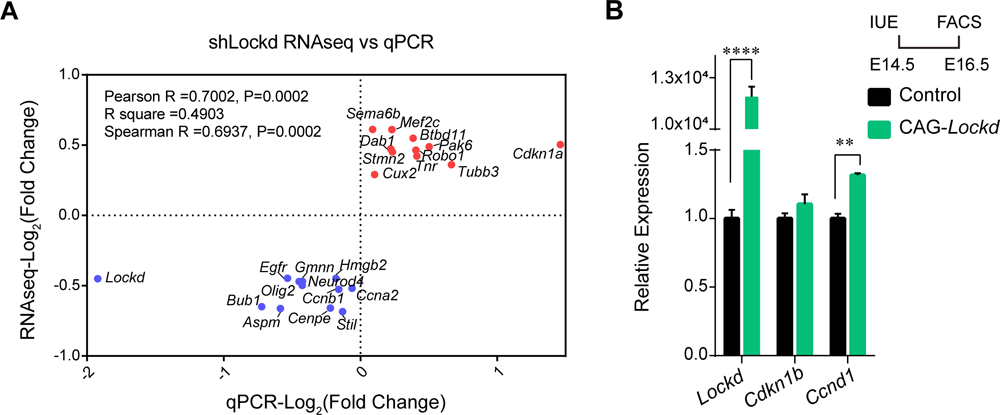
DEGs upon *Lockd* knockdown are related to cell cycle progression in brain development, Related to Figure 5. (A) The mRNA level confirmation of several candidate DEGs identified by RNA-Seq using qRT-PCR in freshly sorted cells normalized to Gapdh. The correlation coefficient was computed between the log_2_ fold change of RNA-seq and qRT-PCR values. (B) The vector expressing full-length *Lockd* driven by CAG promoter and GFP (CAG-*Lockd*-GFP) was electroporated into RGs at E14.5, followed by FACS and RT-qPCR. The empty vector (CAG-GFP) was used as control. Data represent mean ± SEM from technical triplicates (**p < 0.01, ****p < 0.0001; Multiple unpaired t test).

**Figure S5.**
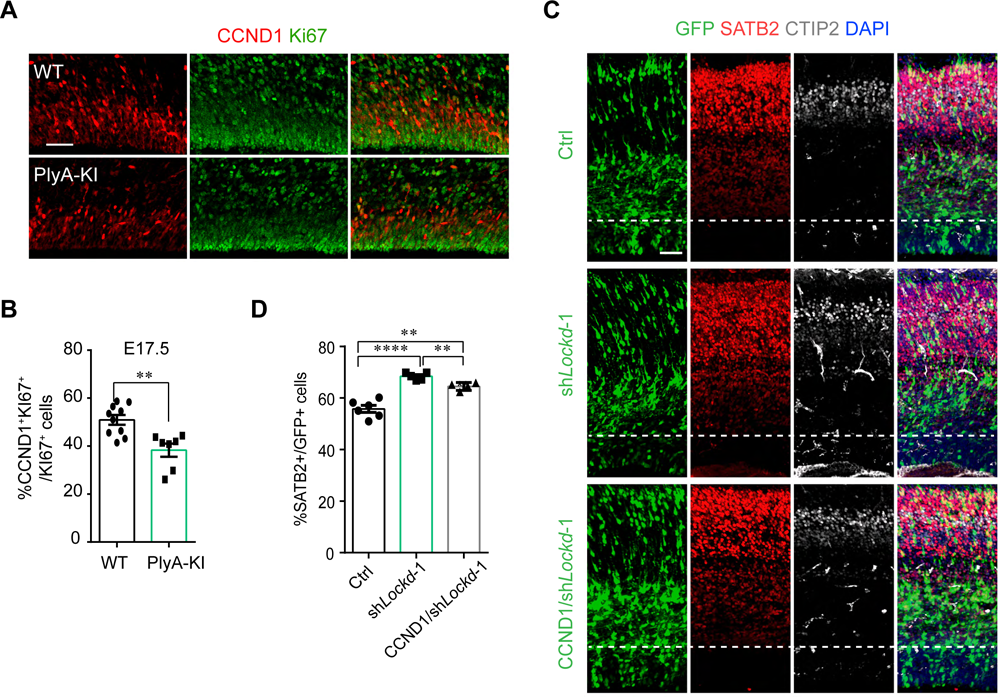
Different *cis-trans* actions of *Lockd* co-regulate the proliferation and differentiation of NSPC, Related to Figure 6. (A and B) E17.5 WT and PlyA-KI cortical sections immunostained for CCND1 and Ki67. The sample confocal images are shown (A; scale bar, 50μm), and the fraction of CCND1^+^/Ki67^+^cells relative to Ki67^+^ cells was quantified (B). Values represent mean ± SEM (n = 10, WT; n =7, PlyA-KI; **p < 0.01; unpaired Student’s t test). (C and D) Neuronal subtypes analysis. E14.5 IUE of sh*Lockd*-1 or coelectroporated with Ef1α-*Ccnd1*, and immunostained with SATB2 and CTIP2 at E17.5 (C, scale bar, 50μm). The percentage of GFP^+^/SATB2^+^ cells among total GFP^+^ cells (D) was quantified. Data represent mean ± SEM; statistical analysis is unpaired Student’s t test (n = 6, Ctrl; n = 5, sh*Lockd*-1; n = 4, CCND1/sh*Lockd*-1; **p < 0.01, ***p < 0.001, ****p < 0.0001).

