## Supplementary material for "Self-balanced regulation by the long non-coding RNA *Lockd* on the cell cycle progression of cortical neural progenitor cells through counteracting *cis* and *trans* roles": Material

| REAGENT or RESOURCE | SOURCE |
| --- | --- |
| **Antibodies** | |
| Mouse monoclonal anti-FLAG M2 antibody | Sigma |
| Rat monoclonal anti-BrdU | Abcam |
| Mouse monoclonal anti-BrdU | Thermo Fisher Scientific |
| Rat monoclonal anti-Ctip2 [25B6] | Abcam |
| Mouse monoclonal anti-SATB2 | Abcam |
| Rabbit monoclonal anti-Ki67 (sp6) | Thermo Fisher Scientific |
| Mouse monoclonal anti-Ki67 | BD Biosciences |
| Rabbit Monoclonal anti-Cyclin D1 (SP4) | Thermo Fisher Scientific |
| Mouse monoclonal anti-Pax6 | Santa Cruz |
| Rabbit monoclonal anti-TBR2 /Eomes | Abcam |
| Sheep polyclonal Anti-Digoxigenin-AP | Roche |
| Chicken polyclonal anti-GFP | Aves Labs |
| Mouse monoclonal anti-Phospho-Histone H3 (Ser10)(6G3) | Cell Signaling Technology |
| Alexa Fluor 546 Donkey anti-Rabbit IgG (H+L) | Thermo Fisher Scientific |
| Alexa Fluor 647 Donkey anti-Rabbit IgG (H+L) | Thermo Fisher Scientific |
| Alexa Fluor 488 Goat anti-Mouse IgG1 | Thermo Fisher Scientific |
| Alexa Fluor 546 Goat anti-Mouse IgG1 | Thermo Fisher Scientific |
| Alexa Fluor 647 Goat anti-Mouse IgG1 | Thermo Fisher Scientific |
| Alexa Flour 488 Goat anti-Rat IgG (H+L) | Thermo Fisher Scientific |
| Alexa Flour 546 Goat anti-Rat IgG (H+L) | Thermo Fisher Scientific |
| Alexa Flour 488 Goat anti-Chicken IgY (H+L) | Thermo Fisher Scientific |
| **Bacterial and Virus Strains** | |
| Escherichia coli (E. coli) | TsingKe |
| Lentivirus | This paper |
| **Chemicals, Peptides, and Recombinant Proteins** | |
| Bromodeoxyuridine (BrdU) | Sigma |
| Paraformaldehyde (PFA) | Sigma |
| DAPI | Invitrogen |
| N-acetyl-L-cysteine (NAC) | Amresco |
| Deoxyribonuclease I from bovine pancreas | Sangon Biotech |
| Fetal Bovine Serum, Qualified, Australia Origin | Thermo Fisher Scientific |
| Poly-L-Lysine (PLL) | Sigma |
| Papain | Worthington |
| N-2 Supplement | Invitrogen |
| B-27 Supplement | Invitrogen |
| Basic Fibroblast Growth Factor (bFGF) | Invitrogen |
| Fluoromount-G | Thermo Fisher Scientific |
| Fastgreen | Sigma |
| TRIzol | Thermo Fisher Scientific |
| Sorbitol | Sigma |
| Deoxyribonuclease (Dnase) | Takara |
| DMEM, High Glucose, no Glutamine | Thermo Fisher Scientific |
| Penicillin-Streptomycin, Liquid | Thermo Fisher Scientific |
| L-Glutamine, 200 mM Solution | Thermo Fisher Scientific |
| Sodium Pyruvate | Sigma |
| Protease Inhibitor Cocktail | Sigma |
| Phosphatase Inhibitor Cocktail 2 | Sigma |
| Bst 3.0 DNA Polymerase | NEB |

| | |
|---|---|
| Deoxynucleotide (dNTP) Solution Set | NEB |
| 50% Dextran Sulfate | Amresco |
| Dulbeccos Phosphate-Buffered (D-PBS) | Sangon Biotech |
| UltraPure™ Formamide | Invitrogen |
| StemPro Accutase Cell Dissociation Reagent | Thermo Fisher Scientific |
| FxCycle™ Violet Stain | Thermo Fisher Scientific |
| **Critical Commercial Assays** | |
| MinElute kit | QIAGEN |
| NucleoBond Xtra Midi EF, for endotoxin-free plasmid DNA | Macherey-Nagel |
| TransScript® One-Step gDNA Removal and cDNA Synthesis SuperMix | TransGen Biotech |
| ChamQTM SYBR qPCR Master Mix | Vazyme |
| Click-iT™ Plus EdU Alexa Fluor™ 647 Flow Cytometry Assay Kit | Thermo Fisher Scientific |
| **Deposited Data** | |
| RNA-seq data | This paper |
| **Experimental Models: Cell Lines** | |
| Neuro-2a | Peking Union Medical College Hospital |
| HEK293FT | Kee. K lab |
| **Experimental Models: Organisms/Strains** | |
| Mouse: ICR | Vital River |
| Mouse: PlyA-KI | This paper |
| **Oligonucleotides** | |
| Primers: in situ hybridization for *Lockd* | |
| F: GCAAGTGCCGTTGACAGAATG | This paper |
| R: GAGATAAGAGATGCAGTCCGGTAG | |
| sh*Lockd* -1: GGTAGAGATGTAGCATCAAAC | This paper, shRNA targeting sequence |
| sh*Lockd* -2: GCCGTTGACAGAATGCAGGCG | This paper, shRNA targeting sequence |
| Primers: pCAG- *Lockd* -hUbc-GFP | |
| F: TTCGAAGCAAGTGCCGTTGACAGAAT | This paper |
| R: GTTTAAACGTGGCTTAAGCCAGATAAAAGTTTATT | |
| Primers: pEF1α -*Cncd1* -IRES-Zsgreen | |
| F: ACTAGTATGGAACACCAGCTCCT | This paper |
| R: GCGGCCGCTCAGATGTCCACATCTCG | |
| siRNA for Control | This study |
| siRNA for *Cdkn1b* | Nguyen et al., 2006 |
| Primers for qPCR | This study |
| **Recombinant DNA** | |
| pEASY-T3 | TransGen |
| p*NeuroD1* -GFP | Guerrier et al., 2009 |
| FUGW-H1- *Lockd* -shRNA1-EGFP | This paper |
| FUGW-H1- *Lockd* -shRNA1-EGFP | This paper |
| FUGW-H1- *Lockd* -shRNA1-TurboRFP | This paper |
| pCAG- *Lockd* -hUbc-GFP | This paper |
| pEF1α -*Ccnd1* -IRES-Zsgreen | This paper |
| **Software and Algorithms** | |
| GraphPad Prism | GraphPad Software |
| Imaris | Bitplane |
| GraphPad Prism | GraphPad Software |
| ZEN | Zeiss |
| Hisat2 v2.1.0 5 | Kim et al., 201 |
| DESeq2 | Love et al., 2014 |

| | |
|---|---|
| Metascape | Zhou et al., 2019 |
| ClicO FS | Cheong et al., 2015 |
| **Other** | |
| Zeiss confocal microscope | Zeiss |
| Electroporator | BEX CO., LTD. |
| Bio-Rad C1000 Thermal Cycler | Bio-Rad |
| Cryostats | Leica |

**IDENTIFIER**

Cat# F1804; RRID:AB_262044
Cat# ab6326; RRID: AB_305426
Cat# B35128; RRID: AB_2536432
Cat# ab18465; RRID:AB_2064130
Cat# ab51502; RRID: AB_882455
Cat# MA5-14520; RRID: AB_10979488
Cat# 550609; RRID: AB_393778
Cat# MA1-39546; RRID: AB_11000217
Cat# sc-81649; RRID: AB_1127044
Cat# ab183991; RRID: AB_2721040
Cat# 11093274910; RRID: AB_514497
Cat# GFP-1020; RRID: AB_10000240
Cat# 9706S; RRID: AB_331748
Cat# A-10040; RRID: AB_2534016
Cat# A-31573; RRID: AB_2536183
Cat# A-21121; RRID: AB_2535764
Cat# A-21123; RRID: AB_253576
Cat# A-21240; RRID: AB_2535809
Cat# A-11006; RRID: AB_2534074
Cat# A-11081; RRID: AB_2534125
Cat# A-11039; RRID: AB_2534096

TSach1-T1 Chemically Competent Cell
N/A

Cat#B5002
Cat#158127
Cat#1612980
Cat#0108-25G
Cat#B100649
Cat#10099141
Cat#P4707-50ml
Cat#LS003126-100 g
Cat#17502048
Cat#17504044
Cat#PHG0261
Cat#0100-01
Cat#F7258-25 g
Cat#15596018
Cat#S8143-1kg
Cat#2212-1000U
Cat#11960044
Cat#15140122
Cat#25030164
Cat#P-5280
Cat#P8340
Cat#P5726
Cat#M0275L

Cat#N0446S
Cat#E516
Cat#E607009
Cat#15515026
Cat#A1110501
Cat#F10347

Cat#28004
Cat#740422.50
Cat#AT311
Cat#Q321
Cat#C10634

GEO: GSE??

N/A
Jung et al., 2017

CD-1(ICR) IGS
N/A

N/A

N/A
N/A

N/A

N/A

Method Details
Method Details
Tables S1

Cat# CT301-01
N/A
N/A
N/A
N/A
N/A
N/A

N/A
N/A
N/A
N/A
https://ccb.jhu.edu/software/hisat2/index.shtml
https://bioconductor.org/packages/release/bioc/html/DESeq2.html

https://metascape.org/gp/index.html
http://clicofs.codoncloud.com

Model: LSM880
CUY21VIVO-SQ
N/A
Model: CM1950

**Primers: Real-time PCR analysis**

| | |
|---|---|
| *Ccnd1-* F | GCGTACCCTGACACCAATCTC |
| *Ccnd1-* R | CTCCTCTTCGCACTTCTGCTC |
| *Polr1a-F* | GCATGGCAGGACGAGAAGG |
| *Polr1a-R* | GATCGTACTGGATGACCAGCC |
| *Cdt1-F* | CGGGCCAAGTCGATCTGTC |
| *Cdt1-R* | ATCCAGGCAGCCGTAACCT |
| *Notch1-F* | GATGGCCTCAATGGGTACAAG |
| *Notch1-R* | TCGTTGTTGTTGATGTCACAGT |
| *Shtn1-F* | CTGGGGACACTTAACAAATCCAC |
| *Shtn1-R* | AGCTTTCTGCGACGTAAAATCC |
| *Nav2-F* | AGTCAAGTCAACAGCAGAGGA |
| *Nav2-R* | CCGGATTTAGCGAGGTAATGATT |
| *Satb2-F* | GCCGTGGGAGGTTTGATGATT |
| *Satb2-R* | ACCAAGACGAACTCAGCGTG |
| *Cplx1-F* | AGTTCGTGATGAAACAAGCCC |
| *Cplx1-R* | TCTTCCTCCTTCTTAGCAGCA |
| *Incenp-F* | AGGCTGAGCGCATGTTTATCA |
| *Incenp-R* | CTCACGGGATCTCTGTTTTCATC |
| *Lockd-F* | GCAAGTGCCGTTGACAGAAT |
| *Lockd-R* | TGAGCAGCCTCCTTTCCAAG |
| *Cdkn1b-F* | TCAAACGTGAGAGTGTCTAACG |
| *Cdkn1b-R* | CCGGGCCGAAGAGATTTCTG |
| *Apold1-F* | CGCTTCCAAGGATTGCTGC |
| *Apold1-R* | CTGAGTGACAACCCCACGAT |
| *Stil-F* | GACACAATTCAGGACTGGTAGAC |
| *Stil-R* | GGCATGATCCACTTTCTGTTCA |
| *Aspm-F* | TGGCTATGAGTGAATGCTCTTCC |
| *Aspm-R* | TCGCGTAAAAACAGTGGCAAG |
| *Cenpe-F* | TCAGGAAAGACACACACGATG |
| *Cenpe-R* | TGCGAGCCATTTCAAAGCCA |
| *Bub1-F* | AGGAGTCAGGTAACAAGGACAT |
| *Bub1-R* | GGACAGTGTCAATGGACGAGA |
| *Ccnb1-F* | AAGGTGCCTGTGTGTGAACC |
| *Ccnb1-R* | GTCAGCCCCATCATCTGCG |
| *Ccna2-F* | GCCTTCACCATTCATGTGGAT |
| *Ccna2-R* | TTGCTGCGGGTAAAGAGACAG |
| *Olig2-F* | TCCCCAGAACCCGATGATCTT |
| *Olig2-R* | CGTGGACGAGGACACAGTC |
| *Neurod4-F* | AGCTGGTCAACACACAATCCT |
| *Neurod4-R* | GTTCCGAGCATTCCATAAGAGC |
| *Hmgb2-F* | GCTCGTTATGACAGGGAGATG |
| *Hmgb2-R* | TTGCCCTTGGCACGGTATG |
| *Gmnn-F* | GGGAGCCCAAGAGAATGTGAA |
| *Gmnn-R* | CAAGCCTTTTGGCAACTCATTT |
| *Sema6b-F* | GCCCTGTCGTTTTTCCTGCT |
| *Sema6b-R* | ACGGGATAGTGGCTCAAGTAG |
| *Dab1-F* | AAAACCAGCGCCAAGAAAGAC |
| *Dab1-R* | CGGACACTTCATCAATCCCAA |
| *Stmn2-F* | GCAATGGCCTACAAGGAAAA |

| | |
|---|---|
| *Stmn2-R* | GGTGGCTTCAAGATCAGCTC |
| *Robo1-F* | GAGCCTGCTCACTTTTACCTC |
| *Robo1-R* | GGTCTGAAGGGTGTTCAACAAT |
| *Btbd11-F* | CCCCAGGGTGATATGAACTCT |
| *Btbd11-R* | AGCAGTTTGCGGAACACATTC |
| *Cdkn1a-F* | CCTGGTGATGTCCGACCTG |
| *Cdkn1a-R* | CCATGAGCGCATCGCAATC |
| *Pak6-F* | CTGTACGCTACTGAGGTGGA |
| *Pak6-R* | GTACCAGCATCCGATCCAGG |
| *Tubb3-F* | TAGACCCCAGCGGCAACTAT |
| *Tubb3-R* | GTTCCAGGTTCCAAGTCCACC |
| *Cux2-F* | GCGGCGTTCCTGAGTGTTTAT |
| *Cux2-R* | CTGGCAGGTGGTTACCGTT |
| *Tnr-F* | GGCTGGAGGTGACTACAGAAA |
| *Tnr-R* | GAAGACCATAGGCTGTTCCTTG |
| *Mef2c-F* | TGTCCAGCCATAACAGTTTGG |
| *Mef2c-R* | CCTTGTGAACATGAAGTCCTCTT |
| *Gapdh-F* | CATGGCCTTCCGTGTTCCTA |
| *Gapdh-R* | CCTGCTTCACCACCTTCTTGAT |
| **SABER-FISH Pool(s) included in** | **Sequence** |
| *Lockd* .27 | AAACTGAAGCTTCCCGCCTGCATTCTGTCAACGGCtttCATCATCAT |
| *Lockd* .27 | TGGACATCCGTGAATGTTTGATGCTACATCTCTACCCtttCATCATCAT |
| *Lockd* .27 | TTTCCAAGACAGGGGAAGGTGCGTCCTCGTCGAACtttCATCATCAT |
| *Lockd* .27 | TGTGAGCAGCCTCCTACGAAAGCACAAATGCAAAAtttCATCATCAT |
| *Lockd* .27 | ACAGGAGACAGAAGGACACAGGGCAAACTTCTTTTtttCATCATCAT |
| *Lockd* .27 | GGTTGCTTCCGCCTTCTCCATTTTGTGGACAGCCAtttCATCATCAT |
| *Lockd* .27 | TGGACTCTCTATCTTTACAGACAAAGAGGACGCCTCAtttCATCATCAT |
| *Lockd* .27 | CCATTCACCAGCTCAAACTCCGTCAAGGACTTTTCtttCATCATCAT |
| *Lockd* .27 | ACAGATCCACCACTGCAGAGCCACTGAACAGGCAGtttCATCATCAT |
| *Lockd* .27 | TTATCTCCTGGTACCCTCCAGGATGCTCCACTCCCtttCATCATCAT |
| *Pax6* .30 | GCTGCTGGGTTGGTGTGTGAGAGCAATTCTCAGATtttAATACTCTC |
| *Pax6* .30 | TAAAGGAGTTGCTCGTGAGAGTTTTCTCCACGGACtttAATACTCTC |
| *Pax6* .30 | TGTTGCTTAAAGACCACAACGGTTTGAAATGACGGTATTTtttAATACTCTC |
| *Pax6* .30 | TCGTTGTCAAGGCTCCTGTCAAGAGTTTTGAGGGCtttAATACTCTC |
| *Pax6* .30 | CGTCCACTTCTAGAATAGAAGATCTCACACATCTGCTCACCtttAATACTCTC |
| *Pax6* .30 | GTTGACAAAGACACCACCAAGCTGATTCACTCCGCtttAATACTCTC |
| *Pax6* .30 | GAGCTAGCTCTACGATCTTCTGCCGGGTGGAGTCCtttAATACTCTC |
| *Pax6* .30 | TCGTAATACCTGCCCAGAATTTTACTCACACAACCGTtttAATACTCTC |
| *Pax6* .30 | CTTACTCCCTCCGATTGCCCTGGGTCTGATGGAGCtttAATACTCTC |
| *Pax6* .30 | ATTTTGCTTACAACTTCTGGAGTCGCCACTCTTGGtttAATACTCTC |
| *Pax6* .30 | CAAAGATGGAAGGGCACTCCCGTTTATACTGGGCTtttAATACTCTC |
| *Pax6* .30 | GCCAGGTTGCGAAGAACTCTGTTTATTGATGACACCtttAATACTCTC |
| *Pax6* .30 | ATACATGCCGTCTGCGCCCATCTGTTGCTTTTCGCTAtttAATACTCTC |
| *Pax6* .30 | GTACTGAAGTCCCGGGATACCAACCAGGGCGTGTGtttAATACTCTC |
| *Pax6* .30 | TCCGTTAGAACTGATGGAGTTGGTGTTCTCTCCCCtttAATACTCTC |
| *Pax6* .30 | AGCTGAAGTCGCATCTGAGCTTCATCCGAGTCTTCtttAATACTCTC |
| *Pax6* .30 | CTTGGGTAAAAGATGTTCTATTTCTTTGCAGCTTCCGCTTCtttAATACTCTC |
| *Pax6* .30 | TAGATCTATTTTGGCTGCTAGTCTTTCCCGGGCAAtttAATACTCTC |
| *Pax6* .30 | ATGTGACTAGGAGTGTTGCTGGCCTGTCTTCTCTGtttAATACTCTC |
| *Pax6* .30 | GGTAGACACTGGTACTGAAGCTGCTGCTGATAGGAtttAATACTCTC |

| | |
|---|---|
| *Pax6* .30 | GTCTGTTCGGCCCAACATGGAACCTGATGTGAAGGtttAATACTCTC |
| *Pax6* .30 | CATGGGTGGCAAAGCACTGTACGTGTTGGTGAGGGtttAATACTCTC |
| *Pax6* .30 | TGCACGAGTATGAGGAGGTCTGACTGGGGACTGGGtttAATACTCTC |
| *Pax6* .30 | ATCATAACTCCGCCCATTCACTGACGGGCTGGTGGtttAATACTCTC |
| *Pax6* .30 | CTGAGACATGTCAGGTTCACTCCCGGGAACTTGGAtttAATACTCTC |
| *Pax6* .30 | TGGCAGAGTGAACACAATTTCCTCTCTCGATCACATtttAATACTCTC |
| *Pax6* .30 | TTCCTGAATACCCAACTGCTGTGTCCACATAGTCATtttAATACTCTC |
| *Pax6* .30 | AGTTACAAAGTGAAGTGCTTCTAACCGCCATTTCTCTTTCTtttAATACTCTC |
| *Pax6* .30 | TGGTTCTAGTCCATTCCCGGGCTCCAGTTCAGGACtttAATACTCTC |
| *Pax6* .30 | CAACTGATACCGTGCCTTCTGTACGCAAAGGTCCTtttAATACTCTC |
| *Pax6* .30 | CGCGATCCAACAGCCTGTGTTGTTCTTTTCTTCATTATAACtttAATACTCTC |
| *Pax6* .30 | TGGAACATCAGTCCATAAACTATGAACAGATGGGGAGAAGTtttAATACTCTC |
| *Pax6* .30 | AGTGTGTGTTGTCCCAGGTTCAATTATATGCAAAGGAATGAtttAATACTCTC |
| *Pax6* .30 | GCTGGCCAAGAAAGAAACAAATGATTGCATGAAAATGTGTtttAATACTCTC |
| *Pax6* .30 | ACAACTGCAAAACACTTAGGTTTAAAACTCTTGCAAGCACTtttAATACTCTC |
| *Pax6* .30 | TGCAGACCTACATACAGCTAGCTGTGTTTTGTTTTAGGTTTtttAATACTCTC |
| *Pax6* .30 | TGCCAGCTCTAATTCAAAACAATTCCTAGTGAATCCCTTGTtttAATACTCTC |
| *Pax6* .30 | GGAATCATTGTGGGGATTGGTTGCCCAGGTTAAAAtttAATACTCTC |
| *Pax6* .30 | GCAACAGCTGTCAGAGGATCTTGTTTAAGAACTCAACCTTTtttAATACTCTC |
| *Pax6* .30 | GGGAGACAGGACAGCAAGAAGGGTTATCTGAGATTGtttAATACTCTC |
| *Pax6* .30 | TTCAATGGGAGCCGATGCCTCTTATGCAAAGAGTGtttAATACTCTC |
| *Pax6* .30 | CACCGAAAGCAAGACACACATGGGAGCAGCTGGAGtttAATACTCTC |
| *Pax6* .30 | CTTGACATGCAACATCCCCTCCACCCTCCAACCACtttAATACTCTC |
| *Pax6* .30 | CCGCTTTGTTGTTGTCTCTTAGTGTGTCTGTGCTCATTATTtttAATACTCTC |
| *Pax6* .30 | CTGAAGGCCTTTAACTCCCACCGGCCAGTCACAGTtttAATACTCTC |
| *Pax6* .30 | CTTCATAGATTTGGGAATGTTTGGCTTAAGCTGCCAATGAtttAATACTCTC |
| *Pax6* .30 | CTTCGCAAATGACAACTGACCAACAATGGGCCCTGtttAATACTCTC |
| *Pax6* .30 | GCATGCTGCCACTTTAAACATGATCAGATCTGTGCTTtttAATACTCTC |
| *Pax6* .30 | TTTTCTCTCCATTGCTTGGGCTCAGGCACTCCTGCtttAATACTCTC |
| *Pax6* .30 | CACAGAACAGGCAGAAAATAGCAGGCAGGCCTTTTtttAATACTCTC |
| *Pax6* .30 | TCAAGCTCGCTGAAGAAAATGTGTCTATGTGACTTAAGTGTtttAATACTCTC |
| *Pax6* .30 | AATCTAGGTGGCCAGGACCCCAGAGTTTCTCTCTCtttAATACTCTC |
| *Tbr2* .26 | CCAGCCATTTCCTCTCGGCTGGCTCTGCTAAACTCtttATAAACCTA |
| *Tbr2* .26 | AGGAAGAAGAGGACTTAGCTCTGAGTTCCCATTTAGCAAtttATAAACCTA |
| *Tbr2* .26 | GCAAGAAAACAAACCCCGGGTTGTAGGTGTCTCTCtttATAAACCTA |
| *Tbr2* .26 | AACAAACTCGGGATTGTAGGTGCCCCTTTCCTTCCtttATAAACCTA |
| *Tbr2* .26 | TAATGTCCCCTTCCTCCCTCGCCTAGGCTACCCACtttATAAACCTA |
| *Tbr2* .26 | AACTGCATGCTTTAGCGAATCGCAGACGGCAACCGtttATAAACCTA |
| *Tbr2* .26 | TAGTAGCGCTCGGAGCTCAGGCTGTCCATGGAGTAGtttATAAACCTA |
| *Tbr2* .26 | AATTTGAGCCATAGGGGCCGGTTGCACAGGTAGACGtttATAAACCTA |
| *Tbr2* .26 | GTTTGGTGATGATCATCTCAGTTTGGTGCCGGTGGtttATAAACCTA |
| *Tbr2* .26 | GGGGTTGAGTCCGTTTATGTTGAAGCTCAAGAAAGGAAAtttATAAACCTA |
| *Tbr2* .26 | AGAACCACTTCCACGAAAACATTGTAGTGGGCGGTtttATAAACCTA |
| *Tbr2* .26 | TGTTATTGTCCGCTTTGCCGCAGGTCACCCACTTGtttATAAACCTA |
| *Tbr2* .26 | GCCTCATCCAGTGGGAGCCAGTGTTAGGAGATTCTtttATAAACCTA |
| *Tbr2* .26 | TGTTGTTTGCACCTTTGTTATTGGTGAGTTTTAACTTCCCAtttATAAACCTA |
| *Tbr2* .26 | CACGATGTGCAGCCTCGGTTGGTATTTGTGCAGAGtttATAAACCTA |
| *Tbr2* .26 | TCATTCAAGTCCTCCACACCGTCCTCTGTCACTTCtttATAAACCTA |
| *Tbr2* .26 | TCTCTGAGAAGGTGAAGGTCTGAGTCTTGGAAGGTtttATAAACCTA |
| *Tbr2* .26 | CGTGTTTTGGTAGGCCGTCACAGCGATGAACTGTGtttATAAACCTA |

| | |
|---|---|
| *Tbr2* .26 | AGCCTTTGGCGAAGGGGTTATGGTCGATCTTTAGCtttATAAACCTA |
| *Tbr2* .26 | ATCTGATGGGATCTAGGGGAATCCGTGGGAGATGGtttATAAACCTA |
| *Tbr2* .26 | AGAAGTTTTGAACGCCGTACCGACCTCCAGGGACAtttATAAACCTA |
| *Tbr2* .26 | TATCGGGCTTGAGGCAAAGTGTTGACAAAGGGCTCCGtttATAAACCTA |
| *Tbr2* .26 | CGTTGGTCTGTGGCACGGTTCTCTCACCGTTATAAtttATAAACCTA |
| *Tbr2* .26 | TTGTTGGTCACAGGTTGCTGGACAGGCGTGACAAGtttATAAACCTA |
| *Tbr2* .26 | GCAAGGACTTAATACCATATGGGAGCAAGGTACTGGAAGTAtttATAAACCTA |
| *Tbr2* .26 | CAGGGTAATACCCCAGGGCATGGGATGTCTGCAGGtttATAAACCTA |
| *Tbr2* .26 | AGTCCAGCTGCCATCTTCCTCTGATAAGCGCCACGtttATAAACCTA |
| *Tbr2* .26 | GGAAGACAGGTGGGCTCATTCTGGATGTCCATGGTtttATAAACCTA |
| *Tbr2* .26 | CGGAGTCGCTGGAGTCTAGAGACTTGATGGAGGGGtttATAAACCTA |
| *Tbr2* .26 | CAGGCGCTTTCTCTTGCAAGCGCTGTTGTACACCCtttATAAACCTA |
| *Tbr2* .26 | TGGAGGTGTCTTTACTGTACTCTTCAGTGTTAATGTCCTCAtttATAAACCTA |
| *Tbr2* .26 | CACAAAACAGGATACATCAAAGGTGGAAGGCAAAAGTCTTTtttATAAACCTA |
| *Tbr2* .26 | GTGGCAAAGCTTTGGCGCCTTCTCTCAGAGAATTGtttATAAACCTA |
| *Tbr2* .26 | ATGCTAGGTCTGCTTCAGTTTAGTTACCTGCGGCCtttATAAACCTA |
| *Tbr2* .26 | AAACACTCCTGCGTCCTCCAGTCACTTTAAAACCAtttATAAACCTA |
| *Tbr2* .26 | AATGAATCAATCCAGCACCTTGAACGACCTTTTCAGTTTTtttATAAACCTA |
| *Tbr2* .26 | TAGCACCGGGCACTCGTTCTTCATAGCCTGCTTTTtttATAAACCTA |
| *Tbr2* .26 | CCCTCTGGGCTGCTGTTTGCTTAAAGATTTATGAGTAGAAAtttATAAACCTA |
| *Tbr2* .26 | ACTCAAGGTCCAACCCTTCTCTTCCCGAAGCAAAAtttATAAACCTA |
| *Tbr2* .26 | CCTGGCAGAAGATATCTGTGAAGAGCCCACTGTTAtttATAAACCTA |
| *Tbr2* .26 | GGAATTGAAGGCAGTGAGTACTAGCTAGCCACTTAAACATCtttATAAACCTA |
| *Tbr2* .26 | GCCAGCCCTACAACAAATGGTTTATTCCAAAATTCTGCAAtttATAAACCTA |
| *Tbr2* .26 | CTCCTGCCTTCTACAGGTCATCCCAGCCACACAGAtttATAAACCTA |
| *Tbr2* .26 | TGGTTTTGCAAATTCTTTACAACGCCAAAGCACCAAATTTtttATAAACCTA |
| *Tbr2* .26 | CGAAGTGGACAGAATATCTCCAAGATATGAATCCCTCCTCTtttATAAACCTA |
| *Tbr2* .26 | GTGTAGAAGCAAGTACGGAGGCAGCTGAGTACTTAGtttATAAACCTA |
| *Tbr2* .26 | GGGGATGACCAAGGAAAGAGGATTAAGCAGGGTTTTtttATAAACCTA |
| *Tbr2* .26 | AGAAGACAGAGCTATACCTGGTCCCTTATTGAACCACATTTtttATAAACCTA |
| **PER reaction for SABER FISH** | |
| Hairpin30.30 | AAATACTCTCGGGCCTTTTGGCCCGAGAGTATTTGAGAGTATT/3InvdT/ |
| Hairpin26.26 | AATAAACCTAGGGCCTTTTGGCCCTAGGTTTATTTAGGTTTAT/3InvdT/ |
| Hairpin28.28 | ACAACTTAACGGGCCTTTTGGCCCGTTAAGTTGTGTTAAGTTGttttttt |
| Hairpin27.27 | ACATCATCATGGGCCTTTTGGCCCATGATGATGTATGATGATGTTTTTTT |
| Branching probe-27*3.28 | ATGATGATGTATGATGATGTATGATGATGTTTCAACTTAAC |
| Clean.G | CCCCGAAAGTGGCCTCGGGCCTTTTGGCCCGAGGCCACTTTCG |
| **Fluor imagers used for SABER FISH** | |
| Fluor imagers-30*.Q670 | /5Quasar670N/ttGAGAGTATTTGAGAGTATTT |
| Fluor imagers-28*.Cy3 | /5Cy3N/ttGTTAAGTTGTGTTAAGTTGT |
| Fluor imagers-26*.Q670 | /5Quasar670N/ttTAGGTTTATTTAGGTTTATT |
